# Compensation for the absence of the catalytically active half of DNA polymerase ε in yeast by positively selected mutations in *CDC28* gene

**DOI:** 10.1101/2020.08.27.269241

**Authors:** Elena I. Stepchenkova, Anna S. Zhuk, Jian Cui, Elena R. Tarakhovskaya, Stephanie R. Barbari, Polina V. Shcherbakova, Dmitrii E. Polev, Roman Fedorov, Eugenia Poliakov, Igor B. Rogozin, Artem G. Lada, Youri I. Pavlov

## Abstract

DNA polymerase ε (pol ε) participates in the leading DNA strand synthesis in eukaryotes. The catalytic subunit of this enzyme, Pol2, is a fusion of two ancestral B-family DNA polymerases. Paradoxically, the catalytically active N-terminal pol is dispensable, and an inactive C-terminal pol is essential for yeast cell viability. Despite extensive studies of strains without the active N-terminal half (mutation *pol2-16*), it is still unclear how they survive and what is the mechanism of rapid recovery of initially miserably growing cells. The reason for the slow progress is in the difficultly of obtaining strains with the defect. We designed a robust method for constructing mutants with only the C-terminal part of Pol2 using allele *pol2rc-ΔN* with optimized codon usage. Colonies bearing *pol2rc-ΔN* appear three times sooner than colonies of *pol2-16* but exhibit similar growth defects: sensitivity to hydroxyurea, chromosomal instability, and an elevated level of spontaneous mutagenesis. UV-induced mutagenesis is partially affected; it is lower only at high doses in some reporters. The analysis of the genomes of *pol2rc-ΔN* isolates revealed the prevalence of nonsynonymous mutations suggesting that the growth recovery was a result of positive selection for better growth fueled by variants produced by the elevated mutation rate. Mutations in the *CDC28* gene, the primary regulator of the cell cycle, were repeatedly found in independent clones. Genetic analysis established that *cdc28* alleles single-handedly improve the growth of *pol2rc-ΔN* strains and suppress sensitivity hydroxyurea. The affected amino acids are located on the Cdc28 molecule’s two surfaces that mediate contacts with cyclins or kinase subunits. Our work establishes the significance of the *CDC28* gene for the resilience of replication and predicts that changes in mammalian homologs of cyclin-dependent kinases may play a role in remastering replication to compensate for the defects in the leading strand synthesis by the dedicated polymerase.

**Author Summary:** The catalytic subunit of the leading strand DNA polymerase ε, Pol2, consists of two halves made of two different ancestral B-family DNA polymerases. Counterintuitively, the catalytically active N-terminal half is dispensable while the inactive C-terminal part is required for viability. The corresponding strains show a severe growth defect, sensitivity to replication inhibitors, chromosomal instability, and elevated spontaneous mutagenesis. Intriguingly, the slow-growing mutant strains rapidly produced fast-growing clones. We discovered that the adaptation to the loss of the catalytic N-terminal part of Pol2 occurs during evolution by positive selection for a better growth fueled by variants produced by elevated mutation rates. Mutations in the cell cycle-dependent kinase gene, *CDC28*, can single-handedly improve the growth of strains lacking the N-terminal part of Pol2. Our study predicts that changes in mammalian homologs of cyclin-dependent kinases may play a role in response to the defects of active leading strand polymerase.

## Introduction

Early studies suggested that two DNA polymerases (pol), pol α, and pol δ, perform replication in eukaryotes [1]. However, the unusual “large” form of pol δ purified from human cells had distinct biochemical properties [2, 3] and appeared to be the third essential eukaryotic pol of the B family, named pol ε [4]. This enzyme has four subunits [5, 6]. Its catalytic subunit (cat-pol ε), encoded by the *POL2* gene in yeast [7], is unusually large and evolved as a fusion of two unrelated ancestral pols of B family [8]. The N-terminal part of cat-pol ε contains a proofreading exonuclease [9] and a catalytically active Fe-S cluster-dependent DNA polymerase [10, 11]. Surprisingly, yeast can survive without this N-terminal pol part, whereas the C-terminal pol part, which lacks both polymerase and exonuclease activities in its evolution [8], is indispensable for yeast growth [12, 13].

Based on the biochemical properties of pol ε, Morrison et al. [7] proposed a model of eukaryotic replication, where pol α and pol δ synthesize the lagging DNA strand and pol ε synthesizes the leading strand. The model proved to be viable, supported by genetic and biochemical experiments, and is currently widely accepted, with some adjustments [14–19]. The main argument against the complete generality of the model was already present in the first paper by Morrison et al., where yeast strains with the insertion of the *URA3* reporter into the middle of the *POL2* open reading frame were viable [7]. This fact remained an annoying puzzle until it was demonstrated that yeast strains with the deletion of the whole N-terminal half of cat-pol ε were viable (mutation *pol2-16*), though grew miserably [12]. Notably, deletions leading to the absence of the whole *POL2* gene or its C-terminal part are lethal [7, 12, 13].

When the active polymerase part of pol ε is deleted, it is likely substituted by pol δ, as proposed by Kesti et al. [12]. The observation that Pol δ can access the 3’OH end of the DNA synthesized by pol ε and proofread leading strand errors [14, 20] supports the feasibility of such pol replacement. The pattern of mutations in strains without the catalytic part of Pol2 is consistent with the participation of pol δ in the synthesis of the leading strand [21, 22]. In contrast to deletion, point mutations impairing the activity of the N-terminal cat-pol ε are lethal [13, 23, 24], supposedly because the presence of the inactivated part does not allow for the substitution by pol δ.

The critical importance of the C-terminal half of cat-pol ε, primarily the C-terminal domain (CTD), was first demonstrated by mutational analysis of the CTD [13]. Later it was found that this domain participates in the formation of a complex with the second subunit of pol ε and the CMG (Cdc45, Mcm2-7, GINS) helicase; thus, its absence prevents the initiation and progression of DNA synthesis [25, 26]. The eukaryotic helicase travels along the leading strand; therefore, pol ε is well-positioned to participate in this strand’s replication [27]. The C-terminal part of cat-pol ε alone is sufficient for vital CMG assembly [28–30], and this role appears to be more critical for replication than the polymerase activity of pol ε. Excessive unwinding by CMG in strains without pol ε activity is mitigated by Rad53 kinase-dependent mechanism [31].

Several groups confirmed and extended the observation that the deletion of the N-terminal part of cat-pol ε in *pol2-16* strains does not stop replication and does not lead to cell death [13, 22, 32]. Strains with the deletion of the N-terminal part of the cat-pol ε grow poorly, are spontaneous mutators, and rapidly acquire the ability to grow at a nearly normal rate. It remained unknown what changes in cells led to the restoration of growth. Besides, several observations in these studies were varying, such as the temperature sensitivity of the corresponding strains [12, 13, 32]. It is likely that the slow progress in finding the mechanism of recovery of cells without the N-terminal cat-pol ε part and discrepancies between different studies resulted from difficulties in creating mutant alleles in the chromosomal location and low stability of truncated protein encoded by *pol2-16* allele [22]. Typically, the *pol2-16* strains without the N-terminal part of Pol2 need two weeks to form visible colonies instead of two days for healthy yeast, and faster-growing isolates might use different adaptation mechanisms [22]. To understand recovery mechanisms, we designed a robust protocol for making better-growing strains with a recoded, codon-optimized galactose-regulatable *pol2rc-ΔN* allele (Materials and Methods) encoding for the C-terminal part of cat-pol ε. The colonies of emerging *pol2rc-ΔN* strains appear three times sooner than colonies of *pol2-16.* The strains with *pol2rc-ΔN* are still slow growers, spontaneous mutators, and frequently yield fast-growing variants. Whole-genome sequencing and genetic analysis of *pol2rc-ΔN* strains provided clues to the nature of genetic events that restore near-normal yeast growth in the absence of the cat part of pol ε. The high proportion of nonsynonymous mutations indicated that the evolution of strains was driven by positive selection and fueled by elevated mutation rates. We show that recurrently occurring single nucleotide changes in the cell-cycle dependent kinase gene, *CDC28*, can singe-handedly restore near-normal growth of strains with the pol ε defect conferred by *pol2rc-ΔN*.

## Results

### Construction of haploid strains without the N-terminal part of cat-pol ε

In previous classic studies, pol mutants were created either by a plasmid shuffling method [23] or by integration-excision into the genome of a vector carrying the allele of interest [9]. The low stability of replicative plasmids is the first method’s limitation, but it is popular because of simplicity [12, 23, 31]. Integration is the most popular method [33–35], but it is not well-suited for examining the phenotypes of lethal mutations or mutations severely reducing fitness. In this method, the mutant copy of a truncated or full-length *pol* allele of interest is integrated into the genome at its original location by transformation by a linearized plasmid. The procedure creates a duplication of the original and plasmid-borne alleles with a selective marker between them. It is essential to use a marker that can be counter-selected, like *URA3*. The single-copy mutant allele is obtained by selecting pop-out with the excision of one gene copy and the *URA3* marker on FOA medium, selective for *ura3* mutants. The integration-excision method works well for haploids and mutator pol alleles [9, 35, 36]. If the mutation leads to lethality or weak growth, the pop-outs will always generate only a wild-type allele reconstituted by recombination. Problematic *pol* alleles could be created in diploids in a heterozygous state, but duplication of a polymerase gene and *URA3* will still be present inside the repeat [24]. Selection for pop-outs in such diploids is difficult (though successes are reported [11, 22]) because the rate of mitotic recombination between the gene and centromere is typically much higher than the rate of intra-chromosomal recombination and thus, most Ura^−^ clones will have a wild-type pol allele. If heterozygous diploids with *polX-URA3-POLX* sporulate, haploid segregants will possess mutant and wild-type alleles, and selection on FOA will again tend to give only clones bearing the wild-type allele. Because of these complications with the integration-excision approach, alternative methods are desirable.

For the reliable construction of haploid strains missing one half of the *POL2* gene, we needed a robust system allowing for the generation of any pol mutants, even those that cause severe growth defects. We used codon-optimized regulatable alleles to alleviate the lower stability of truncated protein encoded by *pol2-16* [22]. We used the following approach to create the *POL2* variant lacking the first half of the gene. At the first step, we created a yeast diploid heterozygous for a complete deletion of the *POL2* gene genetically marked by G418 resistance (**Materials and Methods, Suppl. Fig. 1 and 2**, **Fig. 1A**). Tetrads of the sporulating diploids yielded 2 viable and 2 unviable spores (**Fig. 1B**). All viable spores were sensitive to G418, i.e., possessed a wild-type *POL2* gene. The result is consistent with the known inviability of strains with the deletion of the whole *POL2* gene [7]. In the second step, we integrated either *POL2rc* or *pol2rc-ΔΝ* alleles into the *TRP1* locus of heterozygous diploids using plasmids pJA6 [37] or pJA8 [38], respectively, **Fig. 1A**.

**Figure 1.**
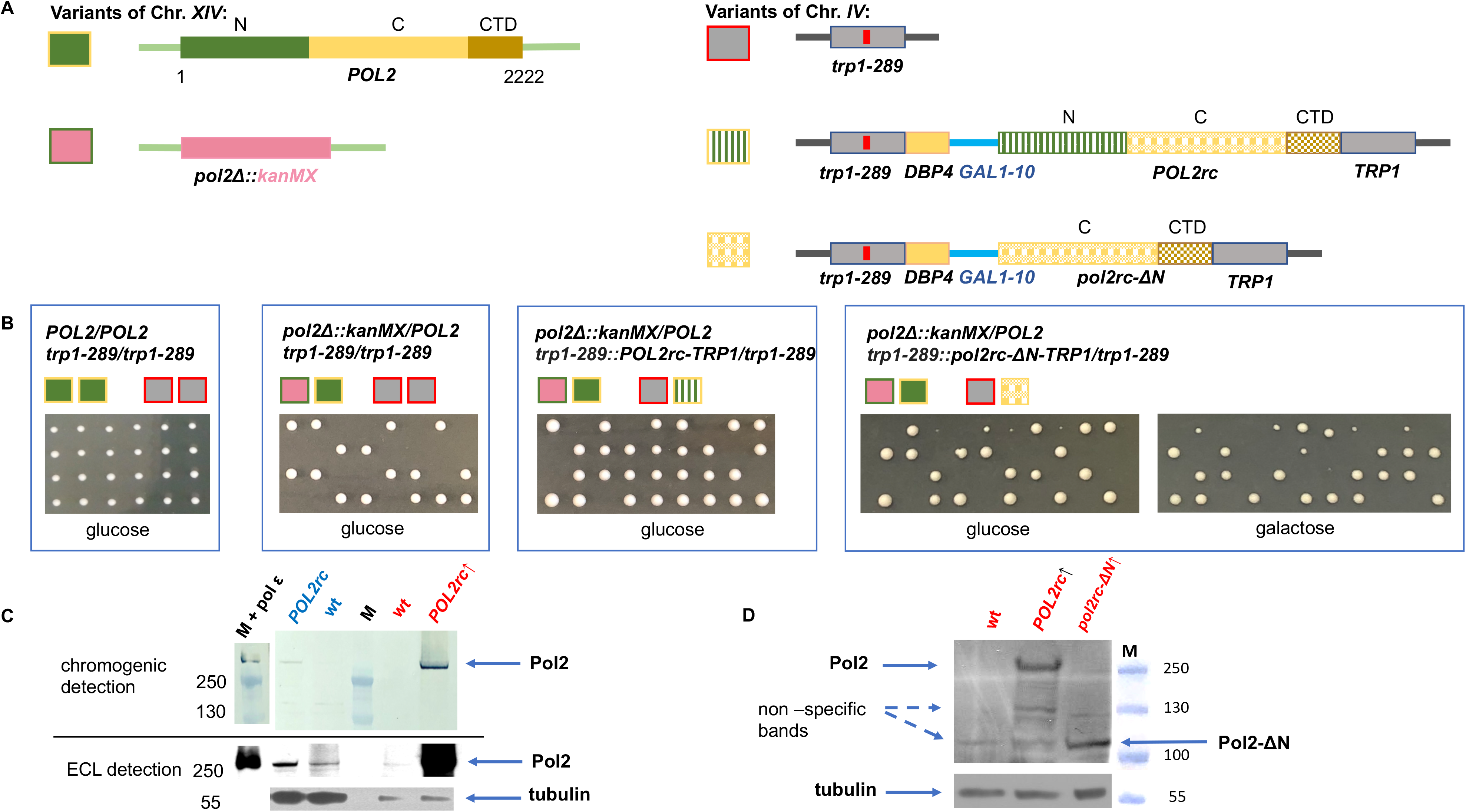
Generation of the *pol2rc-ΔN* haploid strains. **A.** Diploid used for the construction of *pol2rc-ΔN*. The genes in chromosomes XIV and IV are color-coded, and pictograms for each allele are shown to the left of the scheme. The *POL2* gene is diagramed in green (active part) and yellow (inactive C-terminal part). The deletion of the region and substitution with *kanMX* cassette is in pink. The *TRP1* locus is in grey, and the *trp1-289* allele is shown with a red bar inside the ORF representing the mutation. The integration of the optimized *POL2rc* gene into the *TRP1* region is shown by the same colors as the natural *POL2* but with a textured fill. The *pol2rc-ΔN* is in textured yellow. **B.** Results of tetrad dissection of the diploids whose genotypes are represented by pictograms above the photograph. Relevant genotypes of the parent diploids are above pictograms. **C.** Pol2 levels in strain wild-type allele *POL2* (YEE302, *POL2/pol2Δ*) and strain with the recoded *POL2rc* allele (YEE303, *POL2/pol2Δ POL2rc*, Materials and Methods) grown without or without induction of GAL promoter, in YPRAU (raffinose, lanes labeled in blue) or in YPRGAU (raffinose + galactose, lanes labeled in red). Lane labeled “M” is the marker, PageRuler Plus Prestained Protein Ladder covering a range from 10 to 250 kDa (Thermo Fisher Scientific). On lane labeled “M plus pol ε”, in addition to the marker, 2 μL of 2.5 nM purified pol ε were loaded. The upper half represents an image of the membrane developed with the chromogenic substrate (low sensitivity, but the ladder could be seen). The lower part represents the image of the same membrane when the signal is detected by ECL after exposure to film (higher sensitivity). Note that we loaded smaller amounts of extracts prepared from strains grown under induction, in YPRGAU. **D.** Comparable levels of Pol2 (256 kDa) and Pol2-ΔN (110 kDa) proteins in extracts of yeast strain yJF1 (lane 1, labeled wt), or its variants expressing *POL2rc* (lane 2) or *pol2rc-ΔN* allele grown in YPRGAU (lane 3). All five repeats of Western blots gave the comparable ratio of normal and truncated proteins, although different detection kits with a wide range of sensitivity were used (Materials and Methods). Under our conditions and antibodies, truncated Pol2-ΔN protein levels in extracts of strains grown without induction were impossible to determine precisely because of non-specific bands running approximately in the same position.

Tetrad dissection was done on plates with glucose (low levels of expression of *pol* alleles), or on raffinose plates with galactose (high expression of the optimized genes) (**Materials and Methods and Fig. 1B)**. The use of the recoded alleles is advantageous even without induction, even though the western blot signal of Pol2 in extracts without induction in strain with recoded *POL2rc* was just two-fold higher than in *POL2* strains (measured by Image J and adjusted in respect to loading control). Regulatable expression allows for comparable levels of Pol2 to Pol2-ΔN (**Fig. 1D**), which opposes the poor stability of truncated protein seen before in *pol2-16* strains [22]. It is known that *GAL1-10* is leaky and allows for better than natural production of proteins [39], thus, along with the better translation of recoded gene transcripts, explaining the faster appearance of colonies *pol2rc-ΔN* in comparison to *pol2-16.* Genetic segregation of hybrids +/*pol2Δ* heterozygous for *trp-289::POL2rc::TRP1* (insertion of pJA6) followed a typical digenic scenario with one lethal allele and its unlinked suppressor. Most tetrads had three viable spores (tetratype). All viable spores were of the same size, even with no high induction on glucose-containing medium, indicating the full complementation of the deletion by the recoded *POL2rc* allele **(Fig. 1B)** Hybrids +/*Δpol2* heterozygous for *trp-289::pol2rc-ΔN::TRP1* (integration of pJA8) yielded two classes of spores, large and small, all visible on the 4th day of growth. The representative results of the tetrad analysis of diploids heterozygous for the deletion of the natural *POL2* gene and carrying recoded *pol2rc-ΔΝ* (**Suppl. Fig. 1**) are shown in **Fig. 1B (**photo taken on day 4). Small colonies, presumably with *pol2rc-ΔΝ*, grew much better than *pol2-16* strains (natural truncated gene in chromosomal location), especially on galactose-containing medium. The improvement of growth is in stark contrast with the *pol2-16* strains that either did not grow or grew very slowly, forming visible colonies in two weeks [12, 22]. All small colonies were G418-resistant and Trp^+^; thus, the truncated allele *pol2rc-ΔN* was the only source of Pol2 (**Suppl. Fig. 2**). Genome sequencing of these segregants confirmed the normal *POL2* gene’s absence and duplication of the *TRP1* locus (**Suppl. Fig. 3**). In total, we analyzed 53 tetrads of two heterozygous parent diploids, and the overall ratio of tetrads with four viable spores to tetrads with two and three viable spores was close to 1:1:4 (P=0.88) predicted by independent segregation of a wild-type *POL2* on chromosome *XIV* and *pol2-ΔN* on chromosome *IV* (**Suppl. Fig. 1**). Thus, the truncated Pol2-ΔN, lacking the catalytically active part of Pol2, suppresses the lethality of *pol2Δ*. We confirmed by sequencing that *pol2-ΔN* integrated into the chromosome was identical to the original plasmid sequence, ruling out the hypothesis that the truncated inactive pol half acquired polymerase activity by recombination with genes of active DNA pols. We concluded that the elevated expression of the *pol2rc-ΔΝ* allele significantly improves yeast growth, likely providing enough of the C-terminal half of Pol2 for the assembly of the CMG helicase, as compared to the *pol2-16* strains with unstable C-terminal part of Pol2 [21].

### Characterization of pol2rc-ΔN mutants

The analysis of phenotypes of spores with *pol2rc-ΔN* confirmed previous observations made for *pol2-16* strains: they were sensitive to hydroxyurea (HU) (**Fig. 2A**) [12] and slow-growing, rapidly produced faster-growing colonies (**Fig. 2B**) [22]; and had aberrant nuclear and cell morphology in comparison to wild-type yeast (**Fig. 2C**) [22]. Typically, yeast cells are either single or double with smaller buds and larger mother cells. Both types have compact (**Fig. 2C, left panel**). The *pol2rc-ΔN* cells (even from relatively prosperous colonies grown on galactose) show several types of anomalies: large cells, buds with no nuclei, weird-shaped cells, and diffuse nuclei **Fig. 2C**, **right panel).** The proportion of such defective cells reaches 80% in *pol2rc-ΔN* cells in drastic contrast to wild-type cells **(Suppl. Fig. 4).** We did not detect cold-sensitivity (to 20 °C) or thermo-sensitivity (to 37 °C) of *pol2rc-ΔN* strains (**Suppl. Fig. 5),** in contradiction to the earlier study [32]. To understand the reasons for the slower growth of *pol2rc-ΔN* cells, we have used two approaches. In the first, we preferentially stained dead cells with methylene blue. The dye penetrates all cells, but only live cells pump the dye out and remain colorless [40]. We have found that the proportion of methylene blue-stained cells in *pol2rc-ΔN* is 3-5 times higher than in wild-type cells (P<0.01), **Suppl. Fig. 6AB**. The proportion of such cells in galactose is higher in both strains because not every cell easily accommodates to the utilization of galactose, but the trend for the higher proportion of dead cells in *pol2rc-ΔN* is the same. Interestingly, there is no direct correspondence between atypical cells (80%, **Suppl. Fig. 4**) and dead cells (up to 14%, **Suppl. Fig. 6**), suggesting that cell’s repair mechanisms could override the anomalies in pol2rc-ΔN with time. Consistent with that, not all weird-shaped cells are stained by methylene blue (Suppl. Fig 6A). The relatively low number of dead cells only partially explains the slow growth of *pol2rc-ΔN* strains. We obtained additional clues on what happens using a different setup by following individual cells (Materials and Methods), **Suppl. Fig. 7**, **Table 1**. Only a few cells (2%) in wild-type cells do not divide at all in 24 hours, and almost 8% in *pol2rcΔN* strain. The proportion corresponds to the fraction of inviable cells determined by methylene blue staining. Most wild-type cells divide more than three times, while a large proportion of *pol2rcΔN* cells can sustain only one to two divisions for 24 hours **(Table 1).** It is interesting that 76% of *pol2rc-ΔN* cells divide at a rate similar to wild-type cells, forming microcolonies of the same size but with irregular shapes (**Suppl. Fig. 7**, **Table 1).** The median of dividing time of *pol2rc-ΔN* and wild-type cells in the galactose-containing medium is the same five hours in both strains (P=0.341, Mann-Whitney U Test). The results agree with cell cycle analysis of *pol2-16* strains by flow cytometry, where no significant decrease in the velocity of DNA synthesis in *pol2rc-ΔN* cells was detected [12]. The high heterogeneity of *pol2rc-ΔN* cells in the ability to divide adds to the initial slow growth of the colonies and abnormal shape of microcolonies (**Suppl. Fig. 7).**

**Figure 2.**
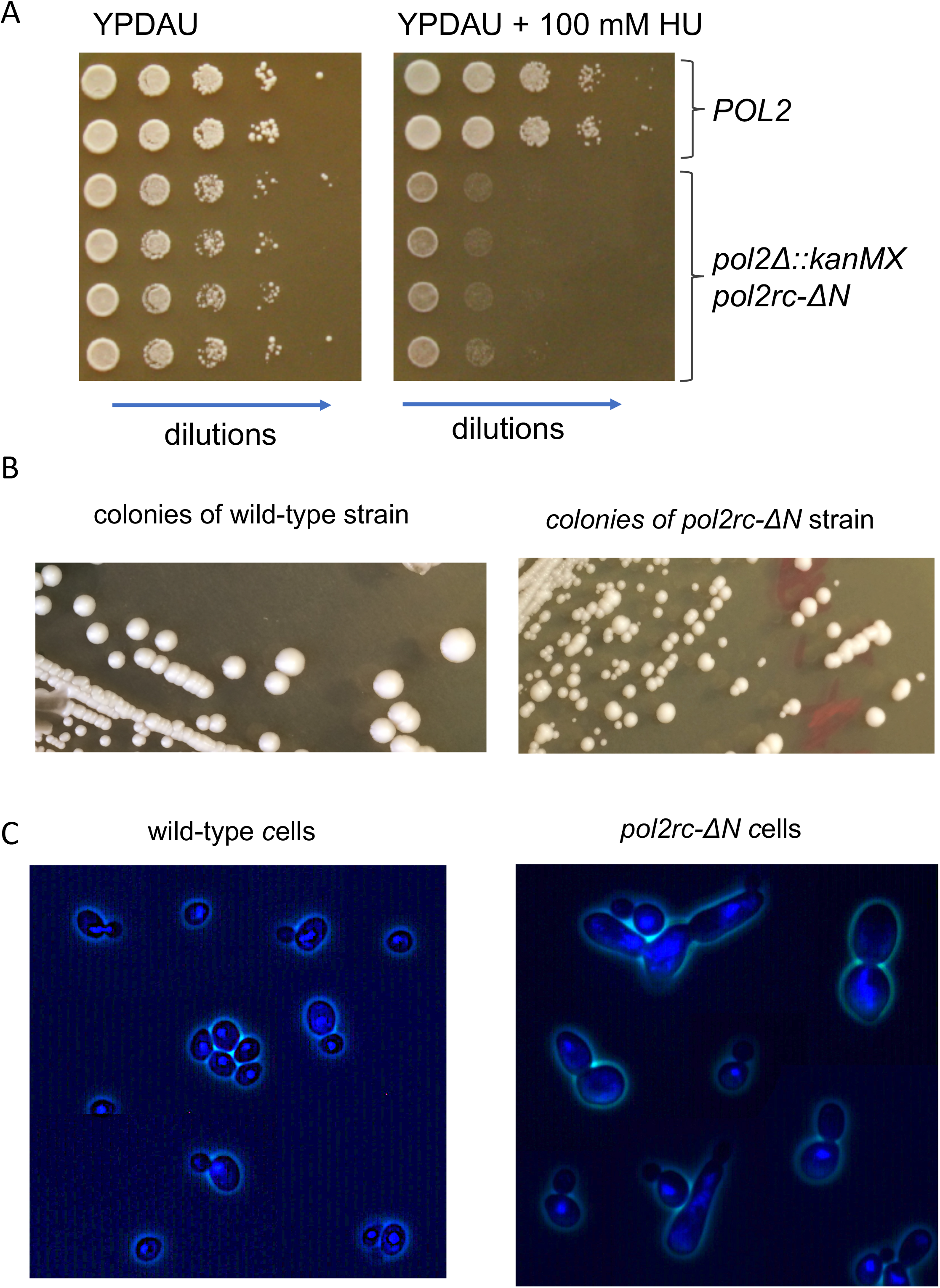
Characteristics of *pol2rc-ΔΝ* strains. **A.** Sensitivity to hydroxyurea (HU) Serial 10-fold dilutions of cell suspension were plated in horizontal rows by a 48-prong device on control or HU-containing YPDAU plates, and photos were taken on day 2. **B.** Small colonies after tetrad dissection (**Fig. 1B**) give large and small colonies when re-streaked onto the YPRGAU medium. The photo on the left is a wild-type strain, and on the right – *pol2rc-ΔN* strain. **C.** DAPI-stained cells from small *pol2rc-ΔN* colonies that were shown in panel **B** in comparison to wild-type cells. The truncation of *POL2* led to aberrant cells with irregular distribution of nuclear material, daughter cells without nuclei, and abnormally large or weird-shaped cells.

**Table 1.**
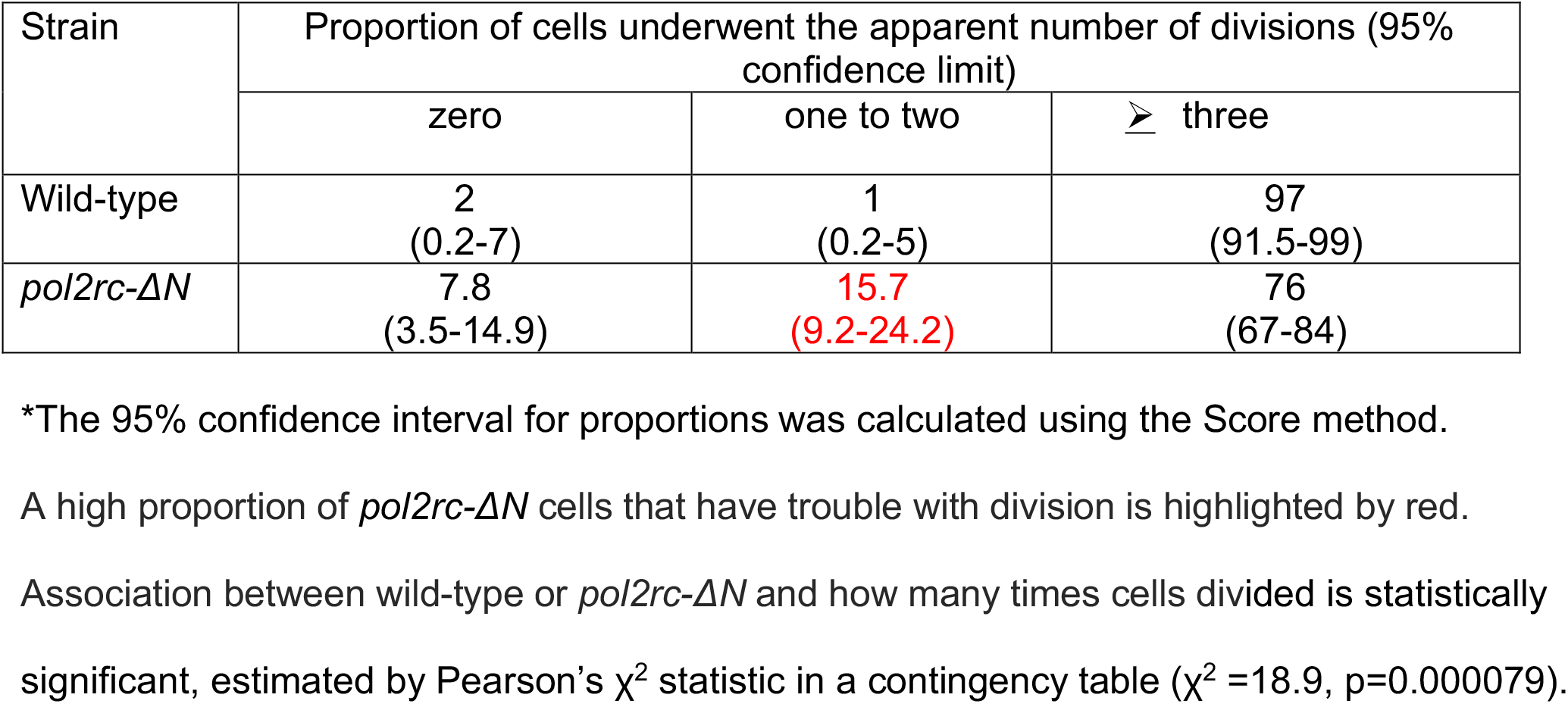
A high proportion of pol2rc-ΔN cells cannot form or have difficulty forming daughter cell buds compared to wild-type cells*.

When we analyzed small colonies grown after tetrad dissection without additional sub-cloning (Materials and Methods, **Suppl. Fig. 2**), segregants with *pol2rc-ΔN* had a 10 to 100 fold elevated spontaneous mutation rate **(Fig. 3),** as was shown previously for the *pol2-1* mutant [41] with the insertion of the *URA3* gene in the middle of *POL2* [7] and the *pol2-16* mutant [21, 32]. The mutator phenotype of *pol2rc-ΔN* strains was highly variable and partially (60%) dependent on the activity of pol ζ (**Suppl. Table 1**), which corresponds to previous observations [21, 32, 41].

**Figure 3.**
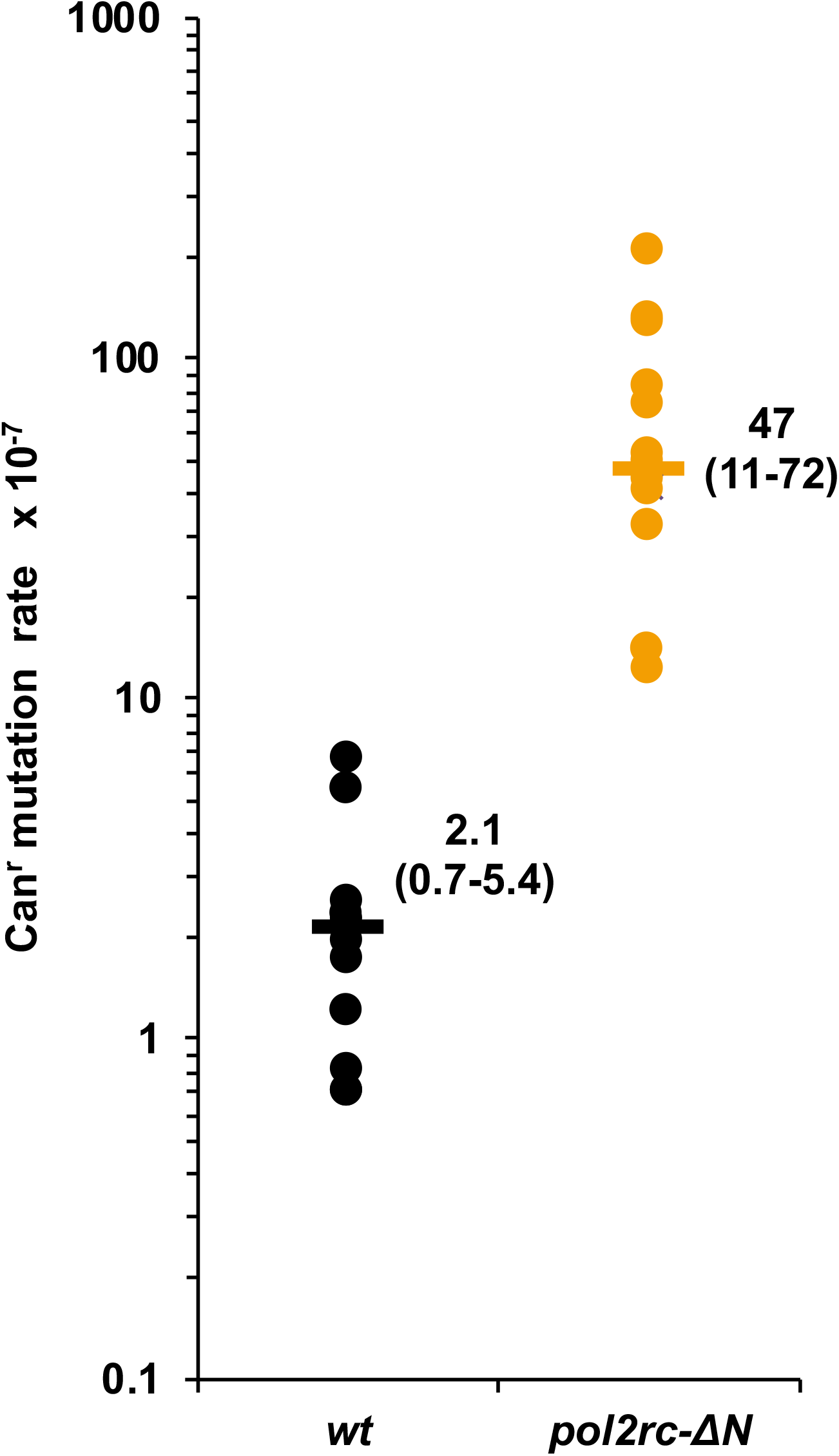
The *pol2rc-ΔN* strains are spontaneous mutators. The rate of spontaneous mutations to canavanine-resistance (Can^r^) is compared in independent *POL2* (black dots) and *pol2rc-ΔN* (yellow dots) segregants. Each dot represents the mutation rate in an independent colony of haploid spore grown on YPRGAU tetrad dissections plates and further cultivated in liquid YPDAU. Horizontal lines show the median values. We present the numerical values of median and 95% confidential interval in the graph’s area in the median’s vicinity.

To learn how the absence of cat-pol ε affects the response to DNA damaging agents, we compared the sensitivity of *pol2rc-ΔN* and wild-type strains to lethal and mutagenic action of UV light (**Fig. 4**). The small difference between the strains in survival was seen only at the highest dose on UV light (**Fig. 4A**, **Suppl.Table 2**). The observation is in partial agreement with the earlier conclusion that *pol2-*16 strains are not sensitive to UV [12]. We found no differences in the induction forward mutations at low and intermediate doses of UV light, and the two-fold decrease at the highest dose (**Fig. 4B)**. We observed a small but significant increase in the frequency of induced reverse frameshift mutations in *pol2-ΔN* strains **(Fig. 4C**, **Suppl. Table 2)**. The result suggests that polymerase activity of Pol ε and the Fe-S cluster located in the N-terminal half of Pol2 are not absolutely required for induced mutagenesis, unlike Fe-S clusters in pol δ and pol ζ [34, 42–46]. Further work is required to find what types of mutations in are attenuated at high doses of UV.

**Figure 4.**
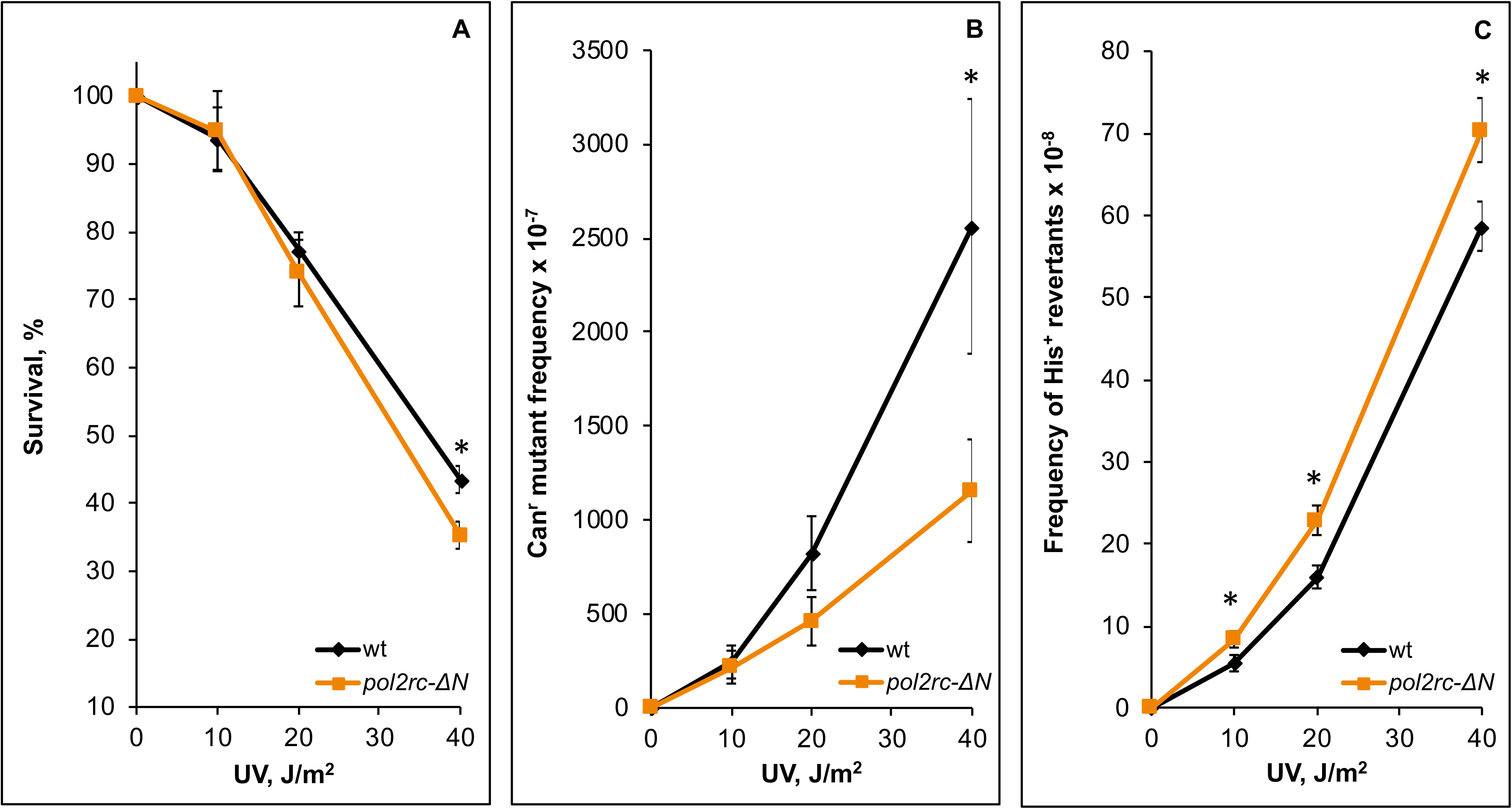
Alterations in UV-survival and mutagenesis in *pol2rc-ΔN* strains. **A**. Survival of *pol2rc-ΔN* (here and in other panels - yellow dots and line) is slightly less than the survival of wild-type strain (here and in other panels - black dots and line) only at the highest UV dose (P=0.0028). Graphs show the mean and standard error of the mean. Data were obtained in two experiments, each with nine independent measurements for every UV dose. A one-tailed t-test was used to evaluate the significance of the difference between means. Asterisks in this and other panels mark statistically significant differences between wild-type and mutant strains. **B**. The frequency of UV-induced forward Can^r^ mutations conferring resistance to L-canavanine in *pol2rc-ΔN* is similar to the wild-type strain, except for a two-fold decrease (P=0.03) at the highest dose. **C.** *The pol2rc-ΔN* strain has an elevated frequency of reversions to His^+^ of the frameshift allele *his7-2* (10 J/m^2^: P=0.02; 20 J/m^2^: P=0.0022; 40 J/m^2^: P=0.01). The mean ± SEM frequency of spontaneous His^+^ reversion is 0.28 (± 0.1–1.2) × 10^−8^ for wild-type clones and 3.9 (± 2.0-7.3) × 10^−8^ for *pol2rc-ΔN* clones.

Severe defects in nuclear morphology in *pol2rc-ΔN* cells indicated difficulties replicating nuclear DNA that may cause chromosomal rearrangements. We used the illegitimate mating test [47] to analyze chromosome stability and revealed that the rate of the loss of chromosome *III* or its arm was elevated 50-fold in *pol2rc-ΔN* strains **(Fig. 5).** The high frequency of chromosomal abnormalities, together with elevated mutation rates, can contribute to the slow growth and cell death in cultures of *pol2rc-ΔN* strains. Point mutations or presumed premutation lesions leading to hybridization of strains of the same “α” mating type were elevated 5-fold, while recombination events leading to hybridizations were not stimulated in *pol2rc-ΔN* strains (**Suppl. Table 3**).

**Figure 5.**
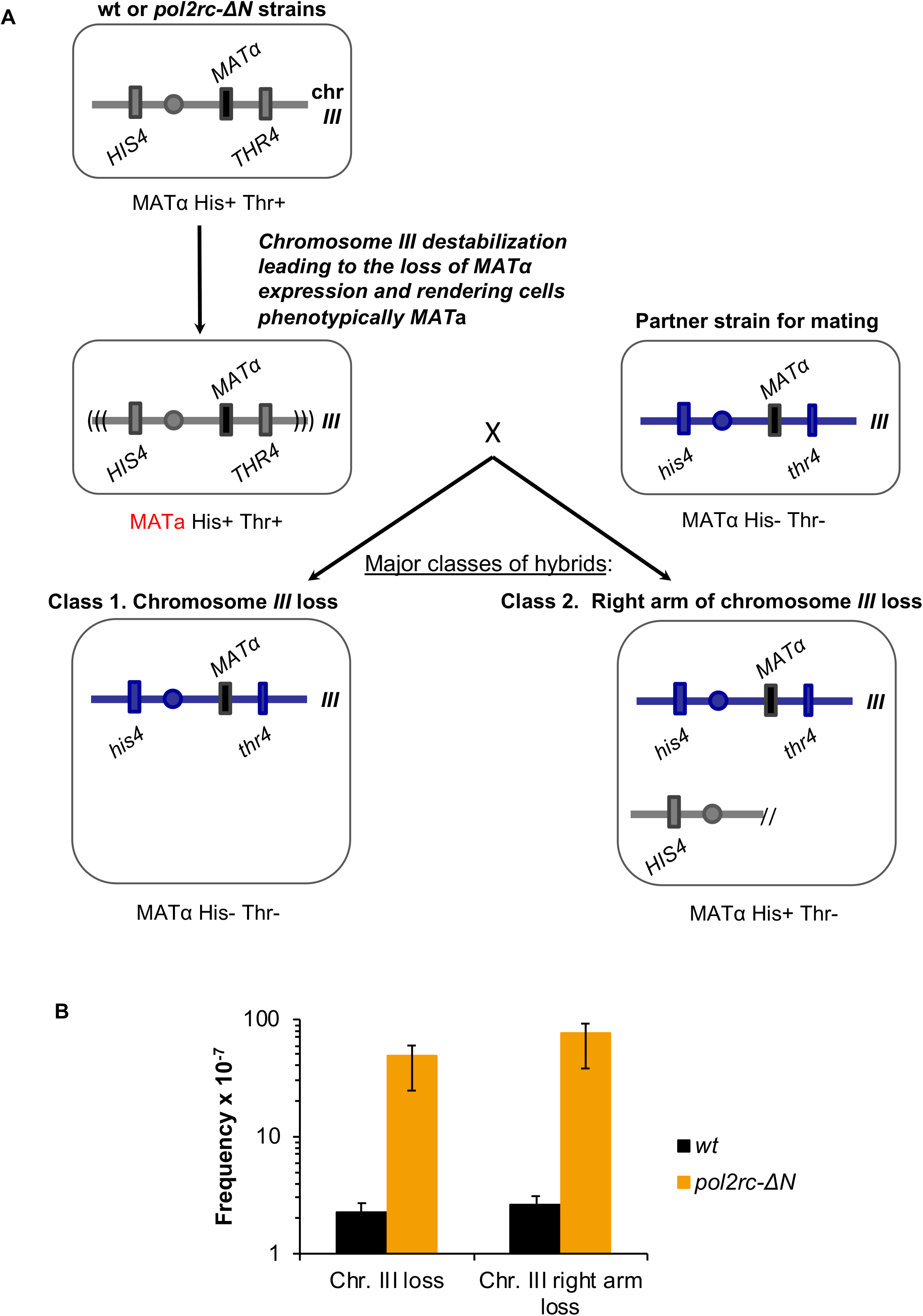
The drastic increase in illegitimate mating of *MAT*α strains in *pol2rc-ΔN* strains. **A.** Schematic of the prevalent genetic events leading to illegitimate mating. In haploid yeast strains, mating type (α or a) is determined by alternative *MATα* or *MATa* alleles of the *MAT* locus. The *MAT* locus is located in the right arm of chromosome *III*. If *MATα* expression is disturbed, heterothallic yeast strains of the mating-type “α” behave like “a” and can hybridize with *MATα* strains [48]. The frequency of this illegitimate hybridization reflects the level of genome destabilization. The mating-type changes in heterothallic strains happen when a cell loses the whole chromosome *III* or its right arm or when the *MATα* locus is mutated. Also, *MATα* may be substituted for by an *HMR**a*** cassette via recombination/conversion mechanism. It is possible to score the frequency of each of these genetic events when strains have selective markers on both arms of chromosome *III*, in our case, *his4* and *thr4* [47, 49]. Phenotype “α” His^−^ Thr^−^ indicates chromosome *III* loss; “α” His^+^ Thr^−^ - for chromosome *III* arm loss. Minor classes (not shown) are non-mater “n/m” His^+^ Thr^−^ and “n/m” His^+^ Thr^+^. They result from recombination/gene conversion between *MAT*α and *HMR***a** and “α” His^+^ Thr^+^ arise by mutations/lesions in the *MATα*. **B.** 50-fold-elevation of the frequency of chromosome *III* or its right arm loss in *pol2rc-ΔN* strains.

### Suppressors of pol2rc-ΔN “slow growth” phenotype

We were intrigued by the fast accumulation of near-normal growing healthy clones in *pol2rc-ΔN* strains (**Fig. 2B**). To detect suppressors enabling strains without the active part of Pol2 to grow, we analyzed genomes of *pol2rc-ΔN* segregants. In the first sequencing run, we analyzed clones of different sizes of four segregants grown on glucose or galactose plates (Materials and Methods). We found approximately the same number of mutations in the genomes of small and large clones (**Suppl. Tables 4 and 5**). When the mutation was present in every genome of all subclones of large and small colonies grown on glucose and galactose (e.g., in *GSP1*, or *RPS1A*, or *OXA1*), it might have happened during the first divisions of the spore after tetrad analysis. Mutations that are present only in genomes of descendants of the large clones are likely to contribute to the improvement of growth. In some cases, new mutations were found only in genomes obtained from the analysis of small colonies consistent with elevated mutation rates in *pol2rc-ΔN* strains. We cannot rule out that at least some of them resulted from the rapid accumulation of fast-growing genetic variants in small clones during their growth for two days in the liquid medium before the isolation of genomic DNA.

Because of no drastic differences in the number of mutations between small and large initial colonies and growth conditions, we next sequenced only two independent clones of 24 *pol2rc-*Δ*N* segregants. The summary of the overall results of genomic sequencing is presented in **Suppl. Tables 4-6,** where the results are presented in several different ways. Overall, we sequenced genomes of 96 strains that originated from 28 independent *pol2rc-ΔN* segregants (Materials and Methods). We have found 292 base pair-substitution mutations. On average, a genome of analyzed strain possessed three new mutations. Three genomes had no mutations, and nine genomes had no base substitutions in coding regions. They may have genome rearrangements that were not detected by our analysis or epigenetic changes. 84 genomes had from one to seven mutations, **Suppl. Table 5**. The same mutations were found in several genomes, **Suppl. Tables 4-6**. The identical mutations found in genomes of common origin (offspring of a single segregant) likely occurred early during spore growth and were recorded as one mutational event. Altogether, we found 197 independent, unique mutations.

We found mutations in genes associated with the cell cycle regulation, DNA metabolism, chromatin, transcription, splicing, translation, cytoskeleton, and other groups **(Fig. 6A).** None of the mutations happened in the *pol2rc-ΔN*, indicating that putative suppressors are all extragenic, quite the opposite to what happens with suppressor od strong mutators *POL2* alleles [50]. Two characteristics of these mutations are relevant to the understanding of their origin. First, the proportion of synonymous changes was less than 8% (**Fig. 6B**), indicating that the appearance of clones with these mutations results from positive selection for mutations improving growth. Typically, in collections of unselected genomic mutations, the proportion of synonymous to nonsynonymous changes does not deviate from a 1:1 ratio by more than 15% [51]. Second, the prevalent type of mutations was changes of CG to GC **(Fig. 6C**, **Suppl. Table 6),** consistent with the participation of pol ζ in mutagenesis in *pol2rc-ΔN* strains (**Suppl. Table 1**). We also found one complex mutation (AAA->TAT), a feature of mutations occurring with pol ζ assistance [52–55]. For most mutations, it is not immediately obvious how certain changes or combinations of changes helped mitigate the growth defect of *pol2rc-ΔN.* The most relevant to the restoration of typical cell division rates are genes that contribute to cell cycle control, DNA repair and recombination, and cell metabolism regulation. Recurring mutations are the most attractive candidates. Among these, independent, nonsynonymous mutations in the *CDC28* gene, encoding for cyclin-dependent kinase, stand out being found five times. In the previous work on the genomic landscape of induced mutations induced by a base analog or APOBEC in yeast, we have found only one mutation (synonymous) among 17,048 mutations [56–58]. Therefore, we focused further on clarifying the role of changes in *CDC28* in the rescue of the growth defect of *pol2rc-ΔN* strains.

**Figure 6.**
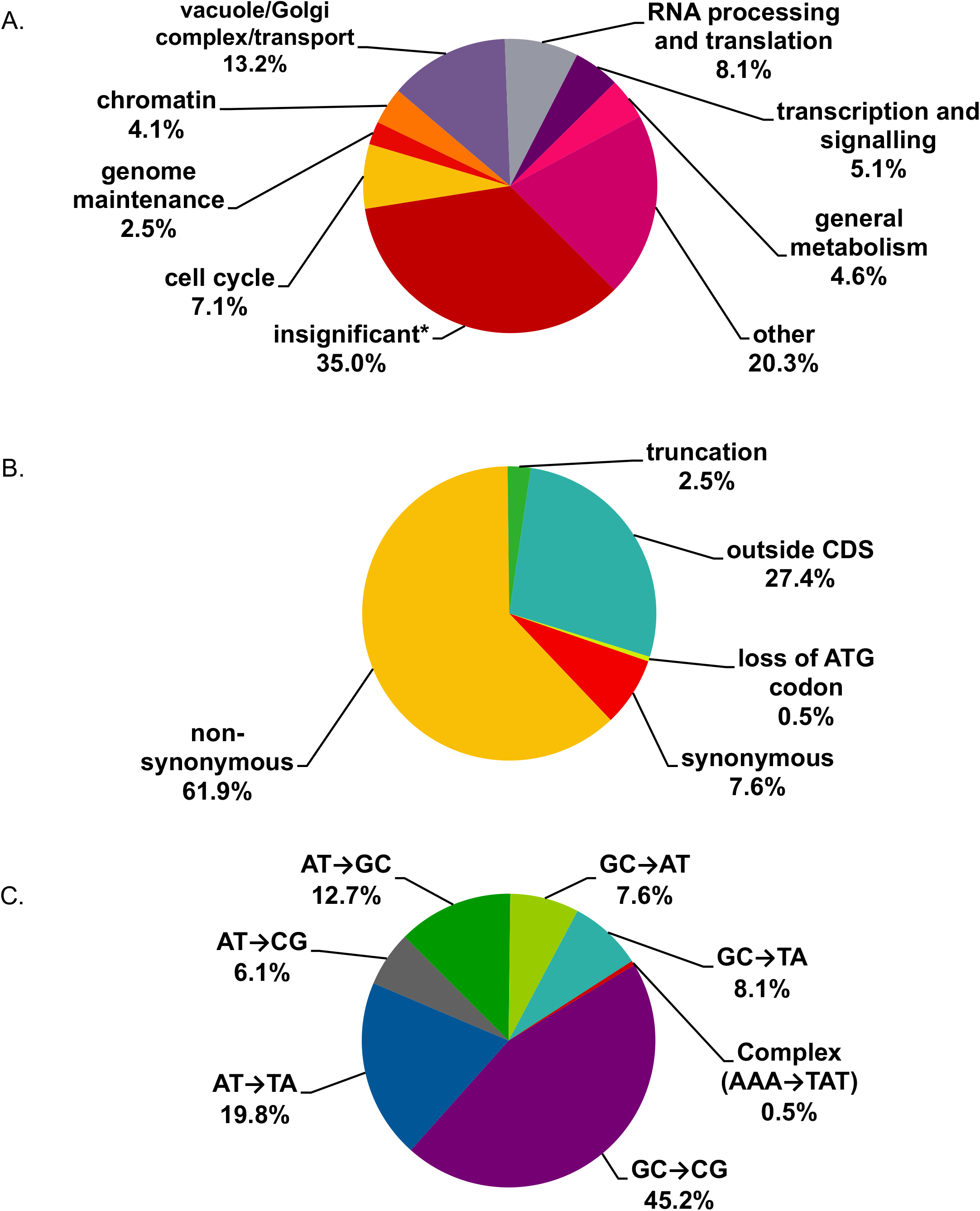
Overview of mutations found in genomes of *pol2rc-ΔN* strains. **A**. Functional groups of genes where mutations were found. * The group named “insignificant” includes synonymous mutations and mutations outside ORFs. **B**. Functional significance of mutations. **C**. Types of mutational changes.

### Mutations in the CDC28 gene suppress the growth defect of the pol2rc-ΔN

Genome sequencing of fast-growing *pol2rc-ΔN* strains (**Fig. 2B**) revealed five independent occurrences of mutations in the *CDC28* gene (**Suppl. Table 4**, **Fig. 7A**). Two independent clones of spore 2A possessed different mutations, *cdc28-186* and *cdc28-257.* The mutation *cdc28-250* was found in clones of two different independent segregants, 59B and 60A. Mutation *cdc28-716* was present in segregant 85C. The probability of getting five independent mutations in the same gene in our collection of sequenced genomes by chance is 1.2 × 10^−6^ (**Supplemental Results 1**). The length of the *CDC28* open reading frame (ORF, 298 codons) is much shorter than the mean ORF length for the yeast genome (491.1 codons; [59], thus the probability value is likely to be underestimated (a conservative estimate of the P-value).To examine the functional significance of *CDC28* mutations and their contribution to the improvement of growth of yeast strains on YPDAU without induction, we crossed each segregant to the wild-type strain LAN201 (described in Materials and Methods, illustrated in **Suppl. Fig. 8,** names of hybrids are in **Fig. 7A**). We reasoned that half of the segregants bearing the *pol2rc-ΔN* allele would get wild-type *CDC28* from LAN201, while the other half would possess the mutated *cdc28* allele (**Suppl. Fig. 8**). The results of the experiments are presented in **Table 2** and illustrated in **Fig. 7B**. Two alleles exhibited almost 100% co-segregation of slow growth with wild-type *CDC28* status and good growth with *cdc28* mutation as determined by the sequencing of G418^r^ Trp^+^ clones. All barely growing colonies (labeled “extra small” in **Table 2**) with *pol2rc-ΔN* possessed the wild-type *CDC28*, and all medium-sized *pol2rc-ΔN* colonies contained either *cdc28-186* or *cdc28-716* (illustrated in **Fig. 7B**). Two mutations, encoding for L84V and L86H, most of the time, with only a few exceptions, segregate with the “good growth” phenotype **(Table 2)**. The effect of these alleles in the rescue *pol2rc-ΔN* is apparent. Additional genetic factors may lead to rare outlier clones (one additional mutation in clone S3 of segregant 2A, three additional mutations in clone SR1 of 59B, and one mutation in SR4 clone of segregant 60A (**Suppl. Table 4)**.

**Figure 7.**
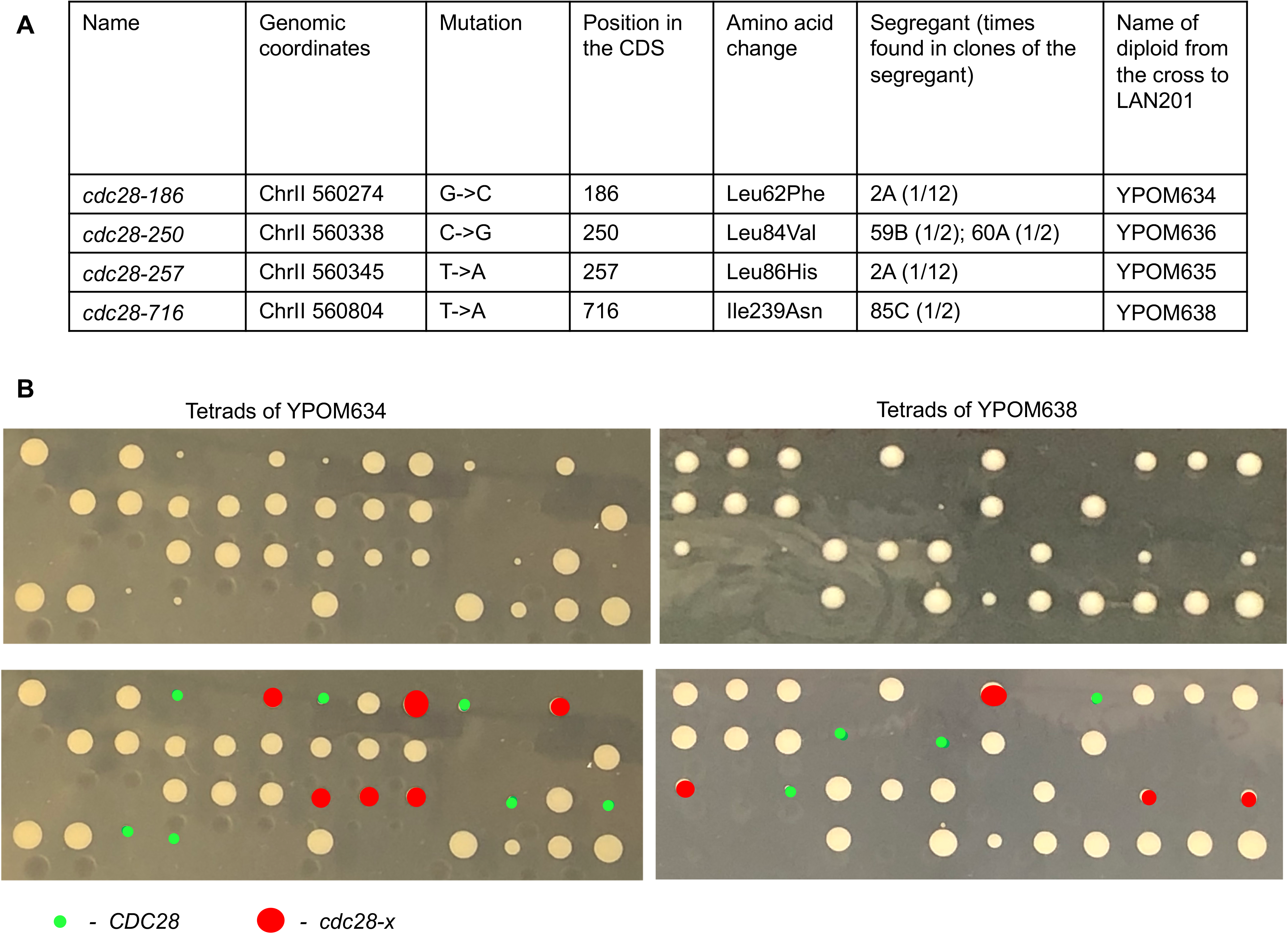
Mutations in *CDC28* suppress the growth defect of *pol2rc-ΔN.* **A**. *cdc28* alleles found during genomic sequencing. **B**. Representative analysis of co-segregation of “good growth” phenotype and alleles of *CDC28.* Upper panel shows the appearance of colonies after tetrad analysis after three days of growth. The lower panel is the same photo where the *CDC28* status of G418^r^ colonies is marked by color.

**Table 2.** The genetic analysis reveals that mutations in *CDC28* efficiently suppress the growth defect of *pol2rc-ΔN* segregants.

| Diploid (*cdc28* allele in heterozygote) | Tetrads dissected | Total viable spores | G418<sup>r</sup> Trp<sup>+</sup> segregants (extra small/medium sized) | Genotyped segregants |  |  |  |
| --- | --- | --- | --- | --- | --- | --- | --- |
|  |  |  |  | Colony size |  | wild-type *CDC28* allele | Mutant *cdc28* allele |
| YPOM634<br>(+/ *cdc28-186*) | 76 | 208 | 71<br>(34/37) | Extra small | 31 | 31 | 0 |
|  |  |  |  | Medium | 32 | 0 | 32 |
| YPOM635<br>(+/ *cdc28-250*) | 60 | 165 | 49<br>(29/20) | Extra small | 23 | 20 | 3 |
|  |  |  |  | Medium | 11 | 1 | 10 |
| YPOM636<br>(+/ *cdc28-257*) | 51 | 143 | 49<br>(23/26) | Extra small | 14 | 13 | 1 |
|  |  |  |  | Medium | 11 | 1 | 10 |
| YPOM638<br>(+/ *cdc28-716*) | 78 | 214 | 67<br>(36/29) | Extra small | 36 | 36 | 0 |
|  |  |  |  | Medium | 26 | 1 | 25 |

We found recurring mutations observed at least twice for future analyses in several other genes (**Suppl. Tables 4, 5**): in *MCM4* encoding for a subunit of CMG helicase [60] in segregants 59B and 73A; in *TRE2*, involved in the regulation of metal transporters [61] in segregants 79B and 82C; in *MNN10*, encoding for a mannosyltransferase [62] in segregants 3B and 15C; in *RPS1A*, encoding for a ribosomal protein [63] in segregants 3B and 19A; and in *BEM2*, encoding for a GTPase activator involved in cytoskeleton organization [64] in segregants 73A and 75D. These mutations are prospects for further investigation. Some mutations were found only once so far, but their occurrence adds to vital the role of *CDC28*-related transactions in the recovery of *pol2-ΔN* strains because they are in the genes encoding for proteins that physically or genetically interact with Cdc28 or regulate cell cycle (e.g., *SIT4, CLB4, CDC5* and *CLB2*, **Suppl. Table 4**).

To approach understanding the mechanisms of suppression, we compared growth on the media panel with different drugs and temperatures of the double *pol2rc-ΔN cdc28-x* strains to single *pol2rc-ΔN*, single *cdc28-x* mutants, and wild-type strains (**Table 3**, **Suppl. Fig 9, Panels A-D**). Mutation *pol2rc-ΔN*, besides sensitivity to HU, leads to sensitivity to free radical producing phleomycin [65] and antimicrotubule drug benomyl [66]. Interestingly, all *cdc28* alleles, when present in *POL2* segregants, do not confer apparent phenotypes associated with alterations of replication: these clones had the same growth parameters as *CDC28* wild-type clones at all conditions examined (compare rows 1 and 7-10 of **Table 3**). All *cdc28* alleles partially rescued HU-sensitivity of *pol2rc-ΔN* at 30 °C (compare row 2 to rows 3-6). In respect to other factors, different *cdc28* alleles showed different properties when combined with *pol2rc-ΔN* (highlighted by red color in **Table 3**). Mutation *cdc28-186* did not suppress the sensitivity of *pol2rc-ΔN* to benomyl (compare rows 2 and 3), *cdc28-250 pol2rc-ΔN* double combination was even more sensitive to phleomycin than single *pol2rc-ΔN* (rows 2 and 4), and *cdc29-716* did not rescue *pol2rc-ΔN* sensitivity to HU at high temperature (rows 2 and 6).

**Table 3.** Mutations in *CDC28* relieve growth defect and sensitivity to cell-damaging agents of *pol2rc-ΔN* segregants*.

| Relevant genotype | 30 °C | 37 °C | HU, 100 mM | HU, 100 mM, 37 °C | Phleomycin, 0.65 μM | UV light, 120 J/m<sup>2</sup> | Benomyl, 90 μM |
| --- | --- | --- | --- | --- | --- | --- | --- |
| wild-type | +++++ | +++++ | +++++ | +++++ | ++++ | +++ | +++ |
| *pol2rc-ΔN* | +++ | +++ | ++ | ++ | ++ | ++ | ++ |
| *pol2rc-ΔN cdc28-186* | ++++ | ++++ | ++++ | ++++ | +++ | +++ | ++ |
| *pol2rc-ΔN cdc28-250* | ++++ | ++++ | ++++ | ++++ | ++ | +++ | +++ |
| *pol2rc-ΔN cdc28-257* | ++++ | ++++ | +++ | +++ | +++ | +++ | +++ |
| *pol2rc-ΔN cdc28-716* | ++++ | ++++ | ++++ | ++ | +++ | +++ | +++ |
| *cdc28-186* | +++++ | +++++ | +++++ | +++++ | ++++ | +++ | +++ |
| *cdc28-250* | +++++ | +++++ | +++++ | +++++ | ++++ | +++ | +++ |
| *cdc28-257* | +++++ | +++++ | +++++ | +++++ | ++++ | +++ | +++ |
| *cdc28-716* | +++++ | +++++ | +++++ | +++++ | ++++ | +++ | +++ |

\*the number of plus signs is an estimate of robustness of growth and corresponds to the number of dilutions where colonies are readily visible. Red font marks the differences in the behavior of the double *pol2rc-ΔN cdc28-x* strains.

## Discussion

Current models of eukaryotic replication assert that the leading DNA strand is synthesized almost entirely by DNA pol ε [17, 67, 68]. The participation of pol α and pol δ in the leading strand was thought to be mostly limited to origins [18, 22] and termination zones [69]. However, Pol δ is unconstrained and can, besides proofreading errors made by pol α [70], proofread errors made by pol ε on leading strand [14, 20] and thus can participate in the synthesis of other parts of the leading strand. Also, Pol δ operates on the leading DNA strand after replication restart when DNA is damaged [71]. Current vision, taking into account all available information, is that pol ε synthesizes more than 80% of the leading DNA strand [68]. Surprisingly, yeast manages to endure the absence of one half of the catalytic subunit of pol ε responsible for DNA synthesis, though growth is poor, at least at first [12]. Other polymerases, most likely pol δ, can substitute for the lost catalytic part [12, 22, 31]. Moreover, cells quickly adapt to the leading strand polymerase’s loss, and colonies start growing almost normally after a relatively short period of struggle ([12, 22], and this study). In the current work, we examined the genetic factors ensuring the replication fork’s resilience and the ability to adapt to the loss of the essential replisome component.

The previous studies could not unequivocally determine the consequence of loss of the N-terminus of Pol2, because the level and stability of C-terminus detected by immunoblotting are much less than of the intact Pol2, and the phenotypes could have resulted from that [22]. Our study overcomes this limitation and allows us to assess the *in vivo* effects of Pol2-ΔN reliably. We utilized a robust method of generation of analog of the *pol2-16* mutation with optimized gene alleles (**Materials and Methods**, **Fig. 1**, **Suppl. Figs 1 and 2**). Expression of the recoded *pol2rc-ΔN* from the galactose-inducible promoter resulted in an improvement of growth not only on galactose-but also on glucose-containing medium **(Fig. 1B).**The level of untagged Pol2-ΔN protein detected by custom-made antibodies against the C-terminal part of Pol2 under induction conditions was similar to that of wild-type Pol2 (**Fig. 1C**). The result suggests that it is not the truncation of Pol2 itself, but the low stability of the C-terminal essential part of the protein [22], that makes *pol2-16* and *pol2rc-ΔN* sick. It appears that when the component of Pol2 necessary for the proper assembly of the CMG complex is abundant, growth is improved. Thus, the C-terminal part (inactive as DNA polymerase) is critical for replication, while the catalytically active N-terminal part is dispensable, as seen before [12], now shown at conditions of better growth of yeast strains with a recoded *pol2rc-ΔN* allele. This observation argues that the 4Fe-4S cluster present in the N-terminal part of Pol2 [10, 11] is not critically involved in the regulation of pol ε’s role in the replisome, though it is needed for the maintenance for the proper architecture of active polymerase. UV-mutability in *pol2rc-ΔN* strains argues that the 4Fe-4S cluster in the catalytic part of pol ε does not play a major role in processes regulating induced mutagenesis, unlike the Fe-S cluster in pol δ and ζ [42, 45, 46, 72]. The difference in UV induced (**Fig. 4**) forward mutagenesis (predominantly base-pair substitutions) and frameshift reversion (+1 and - 2 in poly AT run) [24] is consistent with the different mechanisms of processing of UV damage at regular vs. repeated sequences as suggested by less and variable dependence of UV induced frameshift mutations from pol ζ [73, 74].

The absence of the catalytic function of pol ε is not benign for the cells, even for descendants of the *pol2rc-ΔN* with near-normal growth. The strains are sensitive to hydroxyurea (**Fig. 2A**) and have an increased rate of spontaneous mutation (**Fig. 3**) that partially depends on pol ζ (**Suppl. Table 1**). The latter dependence is not specific to misfunction of pol ε and is characteristic to a broad spectrum of replication defects caused by mutations in genes for the catalytic subunits of all B family pols [24, 41]; accessory subunits [75, 76], parts of CMG complex [77], by high GC content of the templates [78], and replication stress caused by HU [54]. The improvement of growth on galactose-containing medium did not suppress the high level of mutagenesis. Notably, the effect of *REV3* deletion on mutagenesis in *pol2-16* [21] or *pol2rc-ΔN* strains (**Suppl. Table 1**; 40% decrease of mutagenesis) is less pronounced than for other mutations leading to a defective replisome, including the *pol2-1* allele where the *POL2* ORF is disrupted in the middle by the *URA3* marker (>80% reduction of mutagenesis) [41]. The reasons for these peculiarities are still unclear and might reflect the multifaceted role of Pol2 and its catalytically active domain in the replication fork [79]. In addition to the elevation of mutation rate, *pol2rc-ΔN* strains exhibit a high rate of illegitimate mating, resulting from chromosomal rearrangements and losses and enigmatic “primary” DNA lesions temporarily affecting *MAT* locus transcription [47, 49]. These events are signs of replication stress in yeast strains lacking the critical part of Pol2, likely because of the progression of CMG helicase without a corresponding amount of DNA synthesis [31].

The most exciting result came to the fore during the analysis of the remarkable ability of *pol2rc-ΔN* strains to overcome the initial growth defect (**Fig. 2B**). Genome sequencing of independent isolates with this mutation revealed a high prevalence of nonsynonymous changes implying positive selection forces, which, in combination with a mutator effect (with the domination of CG to GC transversions that typically require the involvement of pol ζ [55]), lead to a rapid acquisition of changes allowing for the near-normal growth of *pol2rc-ΔN* strains. Mutations in the genes controlling various pathways, including cell cycle, DNA metabolism, and chromatin remodeling, might have contributed to the effect (**Fig. 6A**), and further analysis will elaborate on the role of these mutations. The most striking case of recurrent mutation has been analyzed in the current work. By genetic analysis, we confirmed that mutations in the *CDC28* gene, the primary regulator of the cell cycle, are connected to the improvement of growth of *pol2rc-ΔN.* Cdc28 in complex with various cyclins inactivates/activates more than 70 cellular targets during cell cycle [80]. One function of Cdc28 is the phosphorylation of the second subunit of pol ε, Dpb2, during the initiation of replication [81]. The complete understanding of how the *CDC28* mutations revive strains without cat-pol ε is a tantalizing task because of the large number of Cdc28 targets and partners [82, 83]. The amino acid residues changed by mutations improving survival of *pol2rc-ΔN* are located on the two different surfaces of the protein, far from the kinase domain (**Fig. 8 A-C**), and most likely modulate interaction with a subunit of the Cdc28 kinase, Csk1/2, with cyclins and other proteins, which, together with Cdc28, regulate cell cycle [65, 84]. Notably, one group of mutations is at the surface of Cdc28 interacting with Clbs, while one mutation affects the surface necessary for interaction with Csk1/2 **(Fig. 8).** In genetic tests, *different CDC28* mutants in combination *pol2rc-ΔN* have individual drug-sensitivity (**Suppl. Fig. 9**, **Table 3).** The plethora of properties of Cdc28 variants found in our study advocates for several ways of suppressing *pol2rc-ΔN* defect. One explanation is based on the observation that the proper level of Cdc28 is required to complete the cell cycle when replication extends beyond chromosome segregation [85]. In the *pol2rc-ΔN* cells, the proportion of under-replicated genomic regions might be higher than in normal cells, as indicated by aberrant cell and nucleus morphology (**Fig. 2C**) and chromosome instability **(Fig. 5B).**The *cdc28* mutations may alter this balance of activity of Cdc28 and other replication factors that results in the better completion of replication in *pol2rc-ΔN* cells. The deficiency of Cks in budding yeast leads to G2 arrest, and elevated levels lead to G2 delay [82]. Of interest, in our collection, there are several mutations in genes encoding those cyclins that participate in the late stages of the cell cycle **(Suppl. Table 4)**.

**Figure 8.**
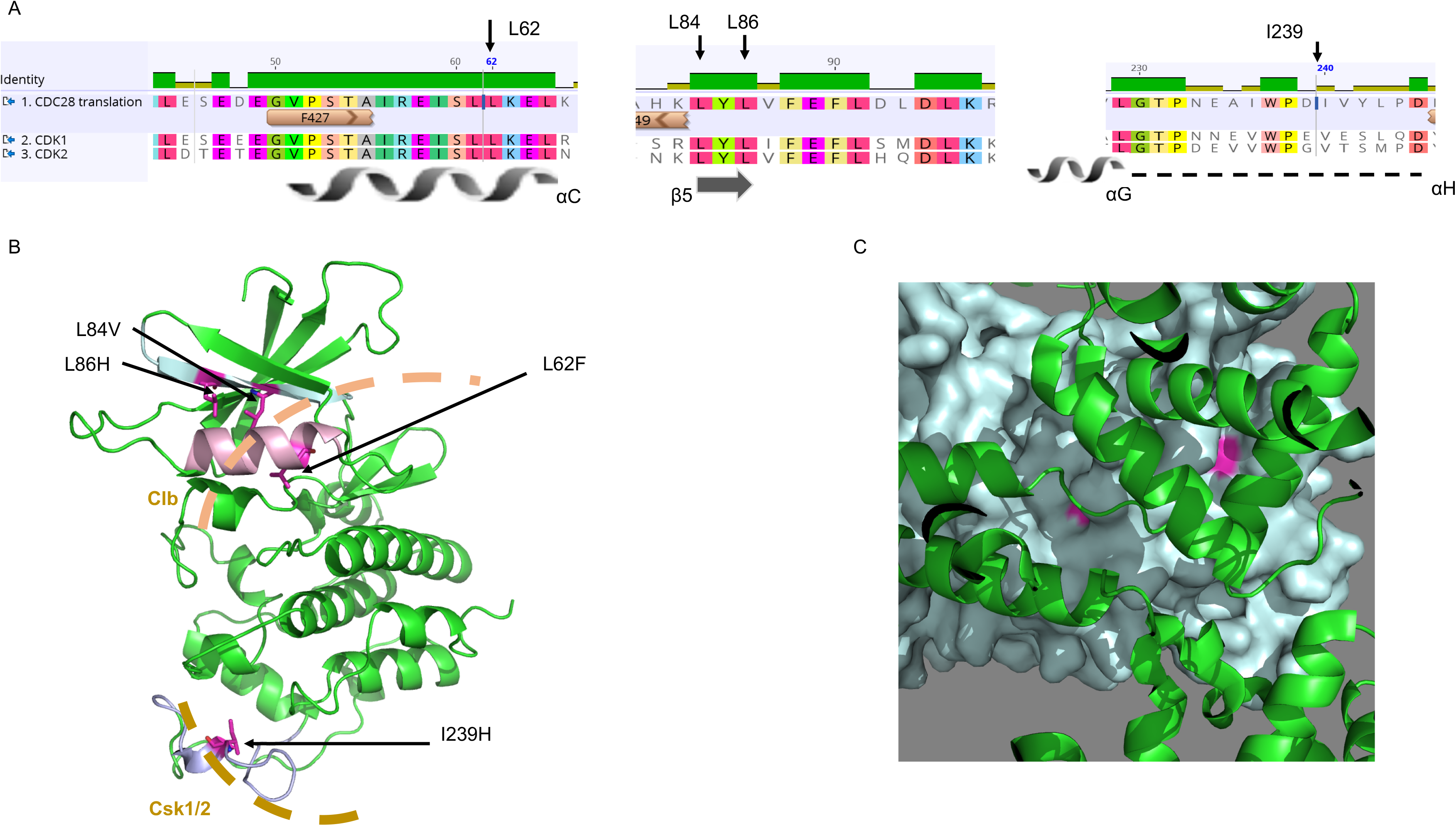
Location of amino acid changes on the modeled structure of Cdc28 in complex with Csk1 and Clb1. A. Amino acid residues that were changed in cdc28 mutants are conserved between yeast Cdc28 and human CDK1 and CDK2. A screenshot of amino acid sequences alignment done by Geneious (Biomatters, Ltd) is shown with secondary structures (α-helical regions as spirals, α-strands as arrows, and loops as dotted lines [83]). B. Amino acid residues that were changed in cdc28 mutants (magenta) are on protein surfaces of modeled Cdc28 (colored green) that interact with Csk1 and 2 (sand-like dashed arc) and cyclins (light brown dashed arc). C. Amino acid residues (pink) that were changed in Cdc28 (teal) at the surface interacting with
cyclin B (green).

Another possibility explaining the effect of alterations in Cdc28 on the recovery of strains without cat-pol ε is that *cdc28* mutations allow for the robust remodeling of the replication fork and faster access of pol δ το the replication of the leading DNA strand. Pol δ can bind the leading DNA strand during replication restart after damage [71]. We can envision that alterations in cell cycle or phosphorylation of DNA pols caused by *cdc28* mutations alleviate pol δ recruitment.

Based on our results, we envisage the following mechanism leading to the appearance of fast-growing clones in *pol2rc-ΔN* strains. The initial defect causes a delay in replication, activation of checkpoint [21] followed by remodeling of the replication fork with the aid of Rad53 kinase, and remaining B-family polymerases replicating both lagging and leading DNA strands [31]. The unusual arrangement of pols leads to increased mutagenesis and genome instability, providing sufficient material for rapid selection against slow-growing cells. It is likely that mutations and, as hypothesized in [21], epigenetic changes may lead to the appearance of fast-growing cells. Changes in many different genes (**Fig. 6A**) can potentially suppress the growth defect of *pol2rc-ΔN*, thus contributing to the ability of cells to adapt to the defect of the regular architecture of leading DNA strand replication machinery. We revealed the prominent role that alterations in Cdc28 play in the resilience of replication machinery.

Our studies in yeast relate to observations on phenotypic manifestation of pol ε dysfunction in higher eukaryotes. In humans, homozygous mutations severely decreasing levels of pol ε are surprisingly not lethal but cause immunodeficiency, facial dysmorphism, livedo, and short stature [86]. Bi-allelic mutations in *POLE* cause IMAGe syndrome [87]. Mutations in *POLE* cause the same conditions as some DNA breakage/instability syndromes [88]. In mice, destabilization of pol ε by a deletion of the gene for the fourth subunit cause growth defects, leukopenia, and a predisposition to cancer [89]. Our work establishing the role of mutations in the *CDC28* in the suppression of growth defects caused by a deficiency of pol ε gene in yeast suggests that variations in its mammalian homologs, *CDK*s, might also play a role in epistatic interactions suppressing defects caused by malfunction of components of pol ε. Multiple alignment of CDK protein kinase orthologs (**Suppl. Fig.10**) suggested that all four amino acid changed in our mutants are located in evolutionarily conserved positions, although the Leu84 site shows some limited variability of residues and instead Ile the 239 site has Val in most species. Mutations in human CDK1 and CDK2, resulting in amino acid changes in the same region that is affected by yeast *cdc28-716*, are found in head and neck and breast invasive ductal carcinoma (CBioPortal, http://www.cbioportal.org/).

## Materials and Methods

### Plasmids

To create strains with recoded galactose-regulatable alleles of *POL2*, we used integrative plasmids with a *TRP1* selectable marker: pJA6, called pRS304(*POL2*+*DPB4-CBP*) with full-length *POL2rc* gene and *DPB4rc* [37]; and pJA8, called pRS304(*POL2* - Δ1261*+DPB4-CBP*) with *pol2rc-ΔN* allele with a nucleotide sequence encoding for amino acid residues 1262-2222 of Pol2 flanked with ATG codon for Met and nonsense codon [38]. The resulting protein is functionally analogous to the protein encoded by *pol2-16* because it similarly misses that N-terminal catalytic active half of Pol2, but is structurally different as *pol2-16* encodes for Pol2 variant with amino acid residues 1-174 fused to 1135-2222 [31, 38]. We named *pol2Δ1– 1261aa* as *pol2rc-ΔΝ*, where “rc” stands for the recoded gene, and ΔΝ stands for the variant of the gene encoding Pol2 without the first 1261 amino acids. All plasmids used in this study were sequenced to verify their identity.

### Yeast strains

We used two related strains, derivatives of CG379 [90]: an autodiploid form of E134 (*MATα ade5-1 lys2::InsE_A14_ trp-289 his7-2 leu2-3,112 ura3-52*) [91, 92], PSD93, described in [39], and an autodiploid of LAN201-Δura3 (*MAT*α *ade5-1 lys2*-Tn*5-13 trp1-289 his7-2 leu2-3,112 ura3-Δ*) [45] described in [93].

Diploid derivative strains heterozygous for a complete deletion of the *POL2* gene (*pol2::kanMX*, **Fig. 1A**, **Suppl. Fig. 1**) were created by one-step gene disruption [94]. At the initial step, a diploid heterozygous for *pol2::kanMX* was constructed by transformation with a PCR product obtained on pFA6-kanMX4 using primers with short homology to the regions flanking the *POL2* gene: pol2_kanMX_45up GAAAGAGCACATTCTATCAAGATAACACTCTCAGGGGACAAGTATCAGCTGAAGCTTCGT ACGC and pol2_kanMX_45downR TTCATGGTAAAGAGGCCATTGAACCTCGCGTTATATACTGCTTACGCATAGGCCACTAGTGGATCTG.

Once the first heterozygous diploid was created and verified, we used its chromosomal DNA as a template for PCR-amplification of the region encompassing *kanMX* with long homology to the region flanking the *POL2:*

POL2 144F TATGGATCTTGATACAGAG

and

POL2 7204R GGCTAATTTTTCGGTTATTT.

These procedures resulted in the generation of PSD93 derivative heterozygous for the deletion allele of *POL2*, named PSD93+/ΔPOL2, and LAN201 derivative named YEE302. The strains were transformed by *Eco*RV-linearized pAJ6 or pJA8 to Trp^+^ to direct the integration of the whole plasmid into the *TRP1* locus, thus creating heterozygotes for the *POL2rc* or *pol2rc-ΔN*, respectively (**Fig. 1A**, **Suppl. Fig. 1**). PSD93+/ΔPOL2 derivatives were named PSD93+/ΔPOL2 [pJA6] or PSD93+/ΔPOL2 [pJA8]. Derivatives of YEE302 with [pJA6] or [pJA8] were named YEE303 and YEE304, respectively. Tetrad dissection of these sporulating diploids gave haploid segregants without the natural *POL2* gene but with recoded gene versions (**Suppl. Fig. 1**). Derivatives of LAN303 and LAN304 strains, heterozygous for *Δrev3*::hph/+ (constructed by one-step gene disruption) were named YEE303r3hΒ/ and YEE304r3hΒ/+, respectively. For analysis of suppressor mutations, we crossed *pol2rc-ΔN* segregants of PSD93+/ΔPOL2 [pJA8] to haploid LAN201-Δura3. Strain 10D-D925 (*MATα ade1Δ his4Δ lys2Δ ura3Δ leu2Δ thr4Δ*) from the strain collection of Department of Genetics and Biotechnology, St. Petersburg University was used in an illegitimate mating assay to score the frequency of whole chromosome *III* and its right arm loss. Original strain for overproduction of variants of pol ε, yJF1 (*MAT**a** leu2-3,112 ura3-1 trp1-1 his3-11,15 ade2-1 can1-100 bar1::hph pep4::kanMX rad5-G535R*) was from J. Diffley laboratory [37, 38].

### Media for yeast

We used standard yeast media [95], but supplemented YPD medium with adenine and uracil for better growth of Ura^−^ and Ade^−^ mutants and prevention of red pigment accumulation in *ade2* strains. Yeast extract and peptone were from “ForMedium™” (Norfolk, UK) at UNMC and from Helicon (Moscow, Russia) and DIA-M (Moscow, Russia). The nomenclature at UNMC is: YPDAU (1% yeast extract, 2% peptone, 2% dextrose, 60 mg/L adenine, 62.5 mg/L uracil); YPGAU (same but with 2% galactose instead of glucose), YPRGAU (same but sugars were 1% raffinose plus 1% galactose); Ccan (synthetic minimal without arginine and with 60 mg/L L-canavanine), HB (minimal with uracil, leucine, threonine, lysine, and histidine). The supplement of adenine and uracil was needed for YPD or YPRG (named here YPD* and YPRG*) in the St-Petersburg laboratory. When necessary, we supplemented media with drugs: 100 mM HU 0.65 μM of phleomycin, or 90 μM of benomyl (all from Sigma-Aldrich, USA).

### Molecular genotyping of yeast strains

We routinely did PCR analysis of integration sites using primers complementary to the junctions of the chromosomal region and vector with recoded genes (the sequences of primers are available upon request). To study the mutations in the *CDC28* gene, we amplified the *CDC28* gene using yeast chromosomal DNA with primers:

CDC28 F242 AGCCAGCACATCAGCTACAGTGG and CDC28 R1414
ACGTCATGGAACACGCCCAGC. The resulting PCR fragment was sequenced with primers: F427 CDC28 GGTGTTCCCAGTACAGCCAT,
F947 CDC28 TGAGATCGATCAGATTTTCAAGA,
R549 CDC28 GCTTGTGTGCATCAGAGTGAA
and R1060 CDC28 GAGGAAAGCTTGGCTTGAAA.

All yeast cultures were grown at 30 °C unless specifically stated.

### Microscopy

Cells were stained by Vectashield Mounting Medium with DAPI reagent according to the manufacturer’s protocol (Vector Laboratories, CA, USA). Imaging of samples was done with a Leica DM 6000B microscope (Leica Microsystems GmBH, Germany) and Leica QWin Standard V3.2.0 software. Fluorescence was detected using the “A” filter component with a 340-nm excitation filter and a 425-nm transmission filter.

### Estimation the proportion of dead and viable cells in yeast cultures

Strains of interest were grown in 200 μL of the appropriate liquid media until the stationary phase (48 hours) at 30 °C. Staining dead cells with methylene blue was done as described before [40], but we used a higher concentration of the dye. Small aliquots of yeas cultures (1-3 μL) were mixed with 30 μL of staining solution (0.2 mg/ml of methylene blue in 2 % sodium citrate) and incubated at room temperature for 5 minutes. 15 μl of yeast suspension was loaded on a microscope slide. At least 1000 cells for each culture were examined under a bright-field microscope (Zeiss Axio Scope.A1), and the percent of dead cells accumulating the blue stain was determined.

### Determination of growth of individual cells

First, we obtained fresh ascospores of the PSD93+/ΔPOL2 [pJA8] strain, using tetrad dissection on complete media with galactose. Then we picked up four wild-type and four *pol2Δ::kanMX trp1-289::pol2rc-ΔN* colonies and inoculated them in liquid YPRGAU media (200 μL). The strains were grown at 30 °C for 48 hours. Then small aliquot of each strain was streaked on the thin agar plates with the same media. Using a microscope and micromanipulator, we picked up 25 healthy cells with no visible buds for each strain and laid them in a square pattern. Cells grown at room temperature were monitored every hour during the next 10 hours to find the time when first, second, and third buds emerged. The data obtained were used to determine the doubling time or the time at which the visible bud had appeared. After 24 hours of incubation, we recorded how many cells produced microcolonies and the final size of the colonies.

### Antibodies for the C-terminal half of yeast Pol2

We used a custom antibody service from ABclonal (Woburn, MA). Two antigenic peptides, STWEVLQYKDSGEPG and AGAWEGTLPRESIV, corresponding to amino acids 1320-1334 and 2182-2195 of Pol2, respectively, were synthesized. Two rabbits were immunized, boosted, and antiserum further purified by affinity chromatography. The company provided purified antibodies at 2 mg/ml concentration.

### Immunoblots

Yeast cells were disrupted by vortexing (2 times 2 min alternated by 1 min ice chill) with 500 μl glass beads in 500 μl PK lysis buffer (50 mM Tris-Cl pH7.6; 50 mM NaCl; 0.1% Triton X-100, 0.1% Tween 20, 1 mM EDTA, 0.5 mM PMSF)[96] or by alkali method [97]. Cell debris and beads were pelleted by centrifugation 14,000g 15 min, and the supernatant was transferred to a new tube. Total protein concentration was determined, then the appropriate amount of extract (50-100 μg) was mixed with loading buffer, denatured, and loaded on 4-12% or 10% NuPage Bis-Tris gel. For control of the sizes of separated proteins, we used PageRuler Plus Prestained Protein Ladder covering a range from 10 to 250 kDa (Thermo Fisher Scientific) and 2.5 nM pure pol ε purified as described [98]. After electrophoresis, proteins were transferred to the PVDF membrane (Millipore). The membrane was incubated for 1 h at room temperature in blocking buffer (5% BSA in 1x TBST buffer) and then cut into sections corresponding to larger and to smaller proteins. The section of the membrane with separated 300 - 95 kDa proteins was incubated with primary antibodies against C-terminus of Pol2 described in the previous section at 1:500 – 1:1,000 dilution in diluent provided in SuperSignal Western Blot Enhancer Kit (Thermo Fisher). Section of the membrane with 75-50 kDa proteins was incubated with rat monoclonal antibodies against yeast α-tubulin (sc-53030, Santa Cruz Biotechnology) at 1:2,000 – 1:4,000 dilution. Incubation with primary antibodies was carried out at 4 °C overnight with gentle rocking. Then, after wash, membranes were incubated in standard blocking solution with appropriate horseradish-peroxidase conjugated secondary antibodies diluted at 1:2000 – 1:4,000: Goat anti-Rabbit IgG, ab97051 (Abcam) or goat anti-rat sc-2006 (Santa Cruz) at room temperature for 1 h. Detection was done by either Western Breeze chromogenic kit or chemiluminescence kit (Pierce™ ECL Western Blotting Substrate, Thermo Fisher) using Kodak BioMax Light film.

### Mutation rates and mutant frequencies determination

The experimental scheme for determination of mutation rate in *TRP1::pol2rc-ΔN pol2Δ* segregants possessing only the C-terminal half of Pol2 is presented in **Suppl. Fig. 2**. Colonies of spores grown on with galactose after tetrad dissection were directly transferred into glucose-containing broth for mutation rates determination. The median rate was calculated by Drake’s formula, as described before, from at least nine spores of the same genotype for each experiment (repeated five times) [24]. UV light mutagenesis was studied as before [45]. Appropriate dilutions of cells were plated on selective and complete plates and irradiated. The mutant frequency in each culture was calculated as a ratio of the number of mutants on selective plates to the number of surviving, colony-forming cells on complete plates. We determined mutant frequency at all UV doses for each culture. These data were used to calculate means and standard error of the mean (Mean ± SEM) from two experiments with nine independent cultures (**Suppl. Table 2**). The UV-induced mutant frequency for each culture was calculated by subtracting the spontaneous background frequency from the mutant frequency in the UV-irradiated cells. Values of UV induced mutant frequency in each culture were used for mean ± SEM determination (**Fig. 4**).

### Illegitimate mating assay

The illegitimate mating assay, or the alpha-test, was used to score the frequency of several distinct genetic changes in chromosome *III*. The principles of the assay were described in detail in [47] [49] and presented in short in the “Results” section. For the assay, aliquots of overnight cultures of nine fresh segregants of *MATα POL2* or *pol2rc-ΔN* strains at appropriate dilution were plated on YPDAU media to estimate the number of live cells in each culture. In parallel, aliquots of the same cultures were plated together with a 10D-D925 strain on HB media for “illegitimate” hybrid selection. After three days of incubation, we counted colonies on HB and YPDAU plates, and then determined the frequency of “illegitimate” hybridization. We tested at least 500 “illegitimate” hybrids for each wild-type and *pol2rc-ΔN* for the mating type and auxotrophy for markers of the left and right arm of the chromosome *III*.

### Genomic DNA sequencing and analysis

We sequenced 96 whole genomes of genotyped segregants (progeny of 28 independent *pol2rc-ΔN* segregants) as well as the parental strains (wild-type and heterozygous for *pol2Δ* variants of PSD93). After tetrad dissection, independent *pol2rc-ΔN* spores were first streaked on glucose-containing YPDAU and YPRGAU media. Among sub-clones of four spores (2A, 3B, 15C, and 19A), we selected 12 colonies for further sequencing: three big and three small colonies from YPDAU media and three big and three small colonies from YPRGAU media. For 24 spores (59B - 90B), we selected only two big colonies from YPDAU media (**Suppl. Fig. 2**). Selected *pol2rc-ΔN* colonies were grown in 10 mL of YPDAU, and chromosomal DNA was isolated by a standard method using phenol extraction [57]. We sheared DNA in M220 Focused-ultrasonicator (Covaris, USA), prepared genomic libraries with the TruSeq® Nano DNA LT Prep Kit (Illumina, CA), and did the whole-genome sequencing using the Illumina HiSeq 2500. We analyzed the raw data using the pipeline described previously [56, 58]. The quality of the raw reads was assessed by FastQC. Reads were filtered and trimmed using Trim Galore with base quality equal to or higher than 20. We mapped reads that passed quality control to our reference strain LAN210_v0.10m using Bowtie2. We updated our pipeline by using tools from the GATK 4.0.3.0 to call (HaplotypeCaller), sort, and filter variants (SelectVariants, VariantFiltration). For the haploid clones, SNVs with AF=0.5 were manually examined in IGV 2.4.10, confirmed to be false positives, and removed from the final vcf file. Full scripts used to generate SNV datasets are available upon request. The raw data are deposited in Sequence Read Archive (SRA, accession # PRJNA634680, Temporary Submission ID: SUB7487615). The BioProject accession number is provided in lieu of SRP and will allow better searching in Entrez.

### Protein structure modeling

The yeast Cdc28 model (P00546) was built on PDB crystal structure 6gu7.2 that was obtained from the Swiss-Model repository. The yeast Cdc28 model was structurally aligned to Human CDK1 in complex with CsK1 and Clb1 (PDB: 5lqf), and mutations in yeast Cdc28 were visualized in the PyMOL software.

## Supporting information

Supplemental results, figures and Tables

## Acknowledgment

We would like to thank John Diffley for sharing plasmids and strains used in the study and Lucy Drury for helping us to understand the details of the constructs. We thank Tahir Tahirov, Andrey Baranovskiy, and Tom Petes for helpful discussions. We thank Elizabeth Moore for excellent technical assistance. We thank Research Park of St. Petersburg State University for next-generation sequencing and UNMC facilities for the sequencing of *CDC28*. The work was done with the financial support of RSF grant # 20-15-00081 and Assignment of Russian Federation Government # 0112-2019-0001 to ES; by the Government of the Russian Federation through the ITMO Fellowship and Professorship Program to AZ; by the National Institutes of Health grants ES015869 and CA239688 to PVS. EP and IBR were supported by the Intramural Research Programs of the National Eye Institute and National Library of Medicine, National Institutes of Health. SRB was supported by the University of Nebraska Medical Center Graduate Studies Fellowship and by the Cancer Biology Training Grant T32CA009476 from the National Cancer Institute. YIP and JC were supported by the Eppley Institute for Research in Cancer Pilot grant. The University of Nebraska Medical Center Genomics Facility and other facilities are administrated through the Office of the Vice-Chancellor for Research and supported by state funds from the Nebraska Research Initiative (NRI), the University of Nebraska Foundation, the Nebraska Banker’s Fund, and by the NIH-NCRR Shared Instrument Program. The facilities also receive partial support from the National Institute for General Medical Science (NIGMS) INBRE - P20 GM103427 and COBRE - P30 GM106397 and P30GM110768 grants, as well as support from the National Cancer Institute (NCI) for The Fred & Pamela Buffett Cancer Center Support Grant-P30 CA036727, The Center for Root and Rhizobiome Innovation (CRRI) 36-5150-2085-20, and the Nebraska Research Initiative. This publication’s contents and interpretations are the sole responsibility of the authors.

## Author contributions

YIP designed the study and wrote the first draft; EIS contributed to the study design, did genetic experiments, and participated in writing on the first draft. ASZ created plasmids and was one of the main contributors to the bioinformatics and statistical part of the work; JC participated in the verification of antibodies, and Western blots, the analysis of mutants; ERT did genetic experiments; SRB participated in strain construction and analysis of proteins; PVS participated in the discussion of work strategy and generated reagents; DEP supervised next-generation sequencing; RF did estimations of the probability of recurrent mutations; EP did molecular modeling and created structural images; IBR supervised the statistical analysis and contributed to the discussion of the significance of the study; AGL led the analysis of the genomic sequences. All authors participated in writing parts of the manuscript and its editing.

## SUPPORTING INFORMATION CAPTIONS

**Supplemental Results 1. Estimation of the probabilities of recurrent mutations in the same gene**

**Supplemental Figure 1. Creation of haploid *pol2rc-ΔN* strains**

**Supplemental Figure 2. Outline of genetic analysis of diploids double heterozygous for the complete deletion of *POL2 gene* and *pol2rc-ΔN***

**Supplemental Figure 3. Peculiarities in the genomic regions of *pol2rc-ΔN* strains relevant to the construction method**

**Supplemental Figure 4. A high proportion of cells with abnormal morphology and nuclear material in***pol2rc-ΔN cells.*

**Supplemental Figure 5. Strains with *pol2rc-ΔN* grow slower than wild-type but are not high temperature-sensitive or cold-sensitive.**

**Supplemental Figure 6. A higher proportion of methylene blue-stained cells in cultures of *pol2rc-ΔN* strains in comparison to wild-type strains.**

**Supplemental Figure 7. Many cells with *pol2rc-ΔN* have difficulty to start dividing**

**Supplemental Figure 8. Schematic of analysis to find if single *cdc28* mutations can confer a growth advantage to *pol2rc-ΔN* strains.**

**Supplementary Figure 9. Rescue of growth defects and sensitivity to drugs of *pol2rc-ΔN* by *cdc28* mutations**.

**Supplementary Figure 10. Multiple alignment of CMGC/CDK/CDC2 protein kinase orthologs.**

**Supplemental Table 1. Mutator effect of *pol2rc-ΔN* is reduced 60% in strain without the catalytic subunit of pol ζ, Rev3.**

**Supplemental Table 2. Differences in mutant frequencies with or without UV-light in *pol2rc-ΔN vs*. wild-type strains.**

**Supplemental Table 3. The *pol2rc-ΔN* elevated chromosome instability measured by the α-test (illegitimate mating).**

**Supplemental Table 4. (Excel file) Summary of mutations found in the course of genomic sequencing of the *pol2rc-ΔN* segregants.**

**Supplementary Table 5. (Excel file) Ancestry of mutations in independent clones of the *pol2rc-ΔN* segregants**.

**Supplemental Table 6. Types of mutations found in genomes of *pol2rc-ΔΝ* strains.**

