## Supplemental results, figures and Tables for "Compensation for the absence of the catalytically active half of DNA polymerase ε in yeast by positively selected mutations in *CDC28* gene": Suppl Fig 1.pdf

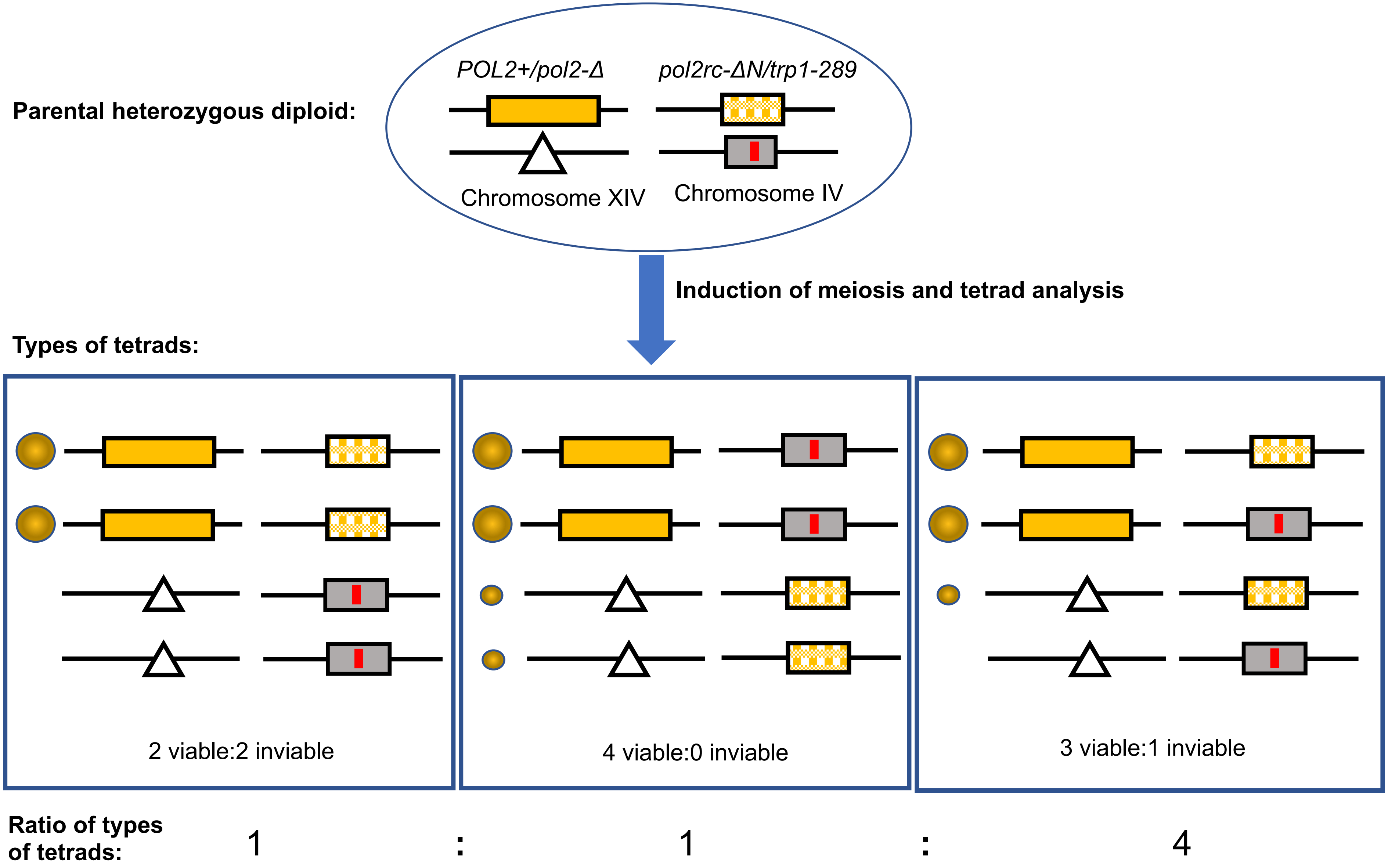


**Supplemental Figure 1. Creation of haploid *pol2rc-ΔN* strains.**
Natural *POL2* gene is colored yellow, recoded truncated gene – yellow with a pattern, shorter then natural allele. Triangle represents deletion of *POL2, trp1-1* allele is represented by a grey box with vertical red line. Each large box represents cartoon of yeast colonies in tetrads to the left and their genotypes to the right. Allele *pol2rc-ΔN is* drawn as a single box to fit into space, but it is actually flanked by *TRP1* and *trp1-289.*
