## Supplemental results, figures and Tables for "Compensation for the absence of the catalytically active half of DNA polymerase ε in yeast by positively selected mutations in *CDC28* gene": Suppl Fig 2.pdf

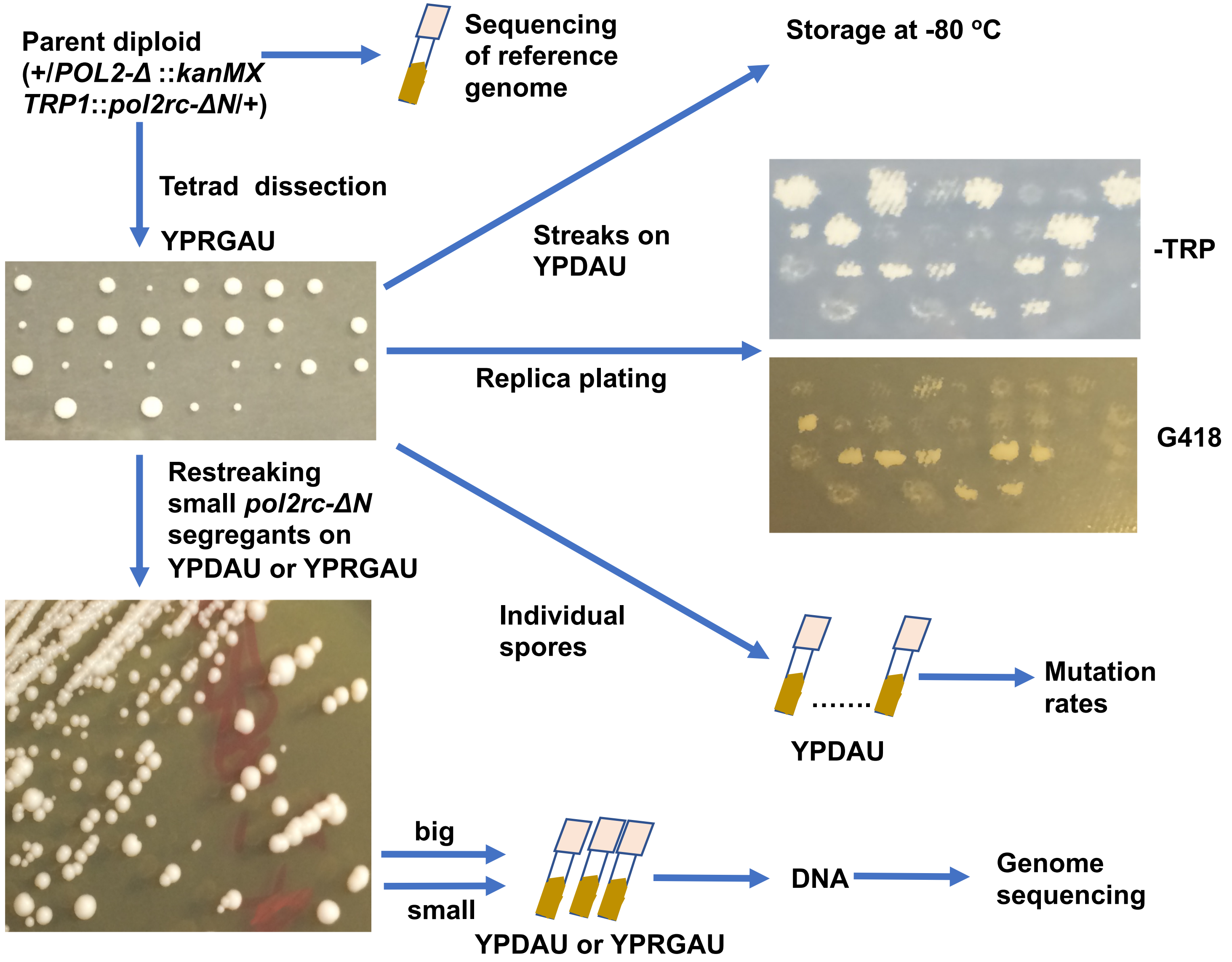


**Supplemental Figure 2. Outline of genetic analysis of diploids double heterozygous for the complete deletion of *POL2* gene and *pol2rc-ΔN*.** Plates with tetrads (upper left) were directly replica-plated to diagnostic media and genotyped segregants were used for mutation rate analysis. Segregants with the combination of the *POL2* deletion and *pol2rc-ΔN* are G418 resistant and $Trp^+$ (upper right). Restreaking of these small colonies gives a plethora of colonies of varying sizes (lower left). These colonies were used for genomic sequencing.
