## Supplemental results, figures and Tables for "Compensation for the absence of the catalytically active half of DNA polymerase ε in yeast by positively selected mutations in *CDC28* gene": Suppl Fig 3.pdf

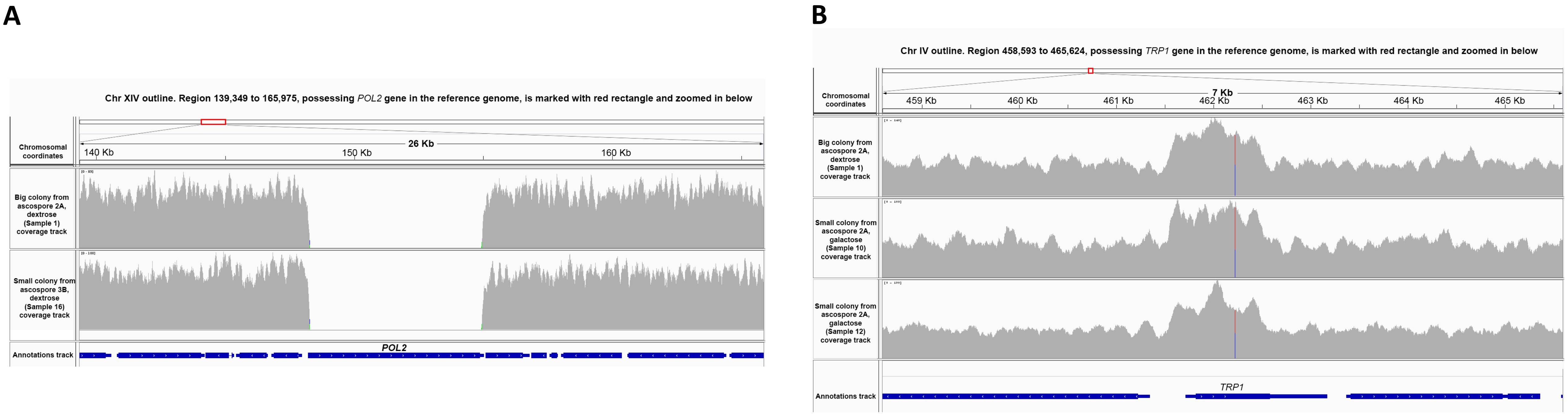


**Supplemental Figure 3. Peculiarities in the genomic regions of *pol2rc-ΔN* strains relevant to the construction method**.
**A**. Genome browser (IGV v. 2.4.10) view of the region of chromosome *XIV* that includes the *POL2* gene in the reference genome LAN210. Sequencing coverage tracks for the chromosomal region (with coordinates on the image) for two sequenced samples (**Suppl. Table 4**) are shown in grey. The annotations track is in blue at the bottom. The absence of the endogenous *POL2* locus is evident because no reads mapped to the corresponding genomic feature (zero coverage).
**B**. Genome browser (IGV v. 2.4.10) view of the region of chromosome *IV* possessing the *TRP1* gene in the reference genome LAN210. Sequencing coverage tracks for the chromosomal region (with coordinates on the image) for three sequenced samples are shown in grey. The annotations track is in blue at the bottom. The coverage is roughly doubled in the *TRP1* genomic feature, indicative of the presence of extra *TRP1* gene copy in the genomes of sequenced clones. The pseudo-heterozygous SNP, shown as a red/blue vertical line on the coverage track, is due to the presence of the *trp1-289* mutation in the reference LAN210 genome, while the wild-type copy of *TRP1* from the integrative plasmid used to insert *pol2rc-ΔN* into the genome is lacking this SNP.
