## Supplemental results, figures and Tables for "Compensation for the absence of the catalytically active half of DNA polymerase ε in yeast by positively selected mutations in *CDC28* gene": Suppl Fig 4.pdf

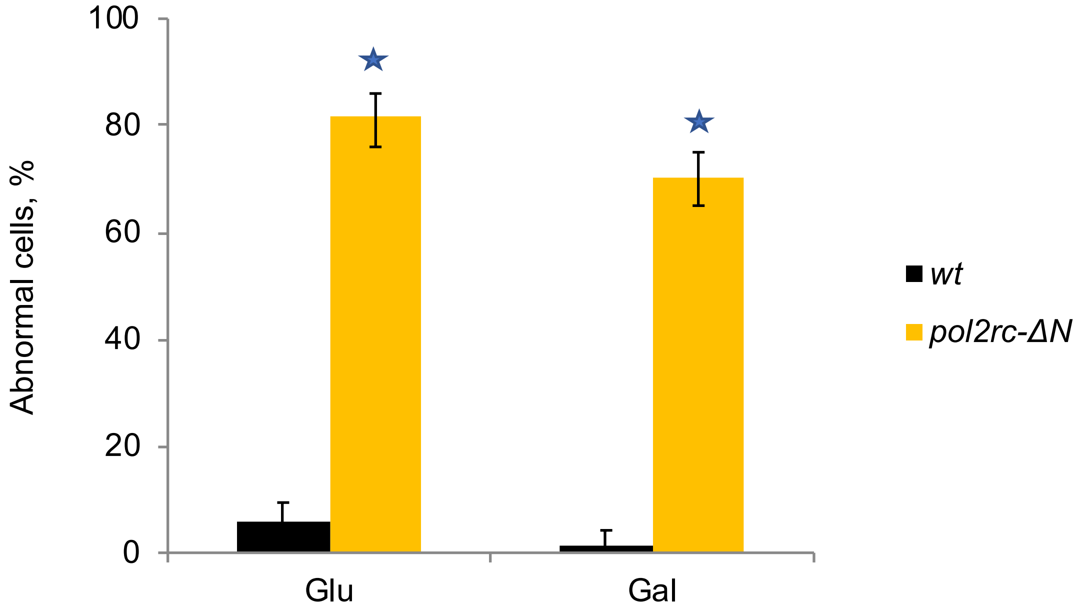


**Supplemental Figure 4. A high proportion of cells with abnormal morphology and diffuse nuclear material in *pol2rc-ΔN cells.***

Glu – YPD*, Gal – YPRG*. For each variant, 200 cells were evaluated. Error bars represent 95% confidence limits.

★- the association between wild-type or *pol2rc-ΔN* and cell's morphology and nuclear status on glucose or galactose media is statistically significant, estimated by Pearson's $\chi^2$ statistic in a contingency table. ($\chi^2$ = 294.4, p= 5.5523E-66 and $\chi^2$ = 230.9, p=3.7889E-52, respectively for glucose or galactose media).
