## Supplemental results, figures and Tables for "Compensation for the absence of the catalytically active half of DNA polymerase ε in yeast by positively selected mutations in *CDC28* gene": Suppl Fig 5.pdf

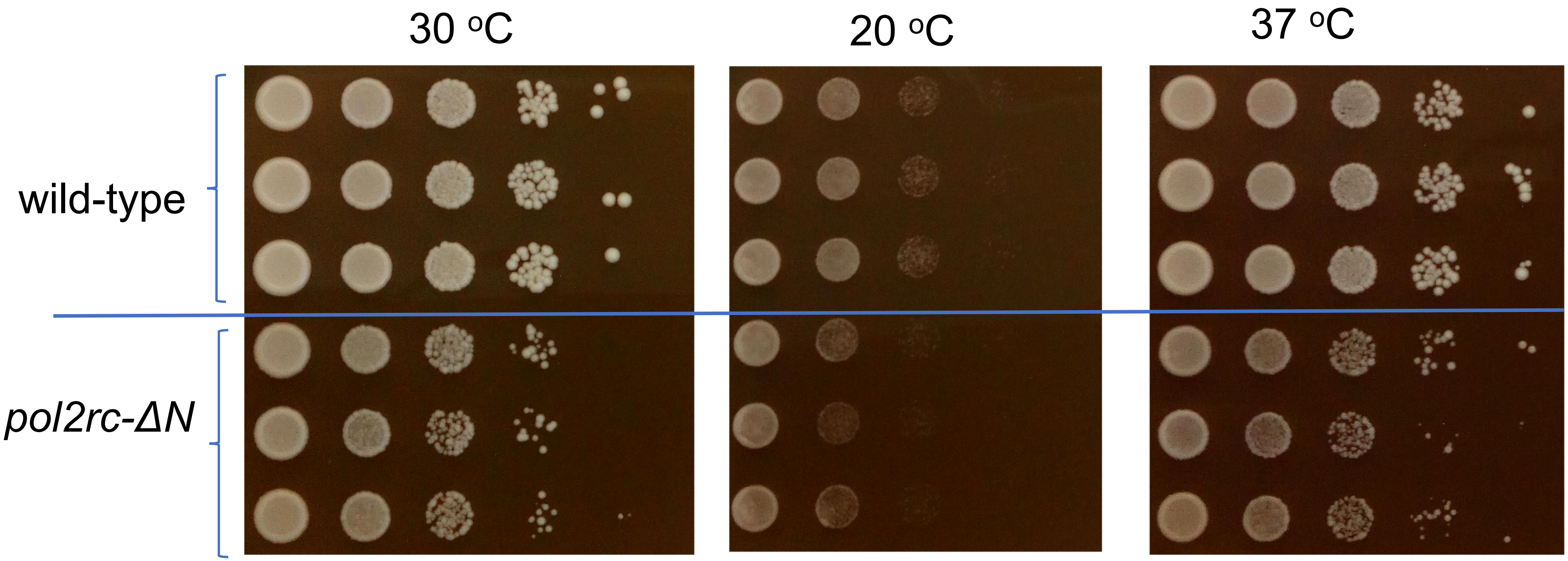


**Supplemental Figure 5. Strains with *pol2rc-ΔN* grow slower than wild-type but are not high temperature-sensitive or cold-sensitive.**

Serial 10-fold dilutions of cell suspension of strains grown in YPRGAU were plated in horizontal rows by a 48-prong device on control, precooled/preheated plates with YPDAU media and incubated at the indicated temperature for two days.
