## Supplemental results, figures and Tables for "Compensation for the absence of the catalytically active half of DNA polymerase ε in yeast by positively selected mutations in *CDC28* gene": Suppl Fig 6.pdf

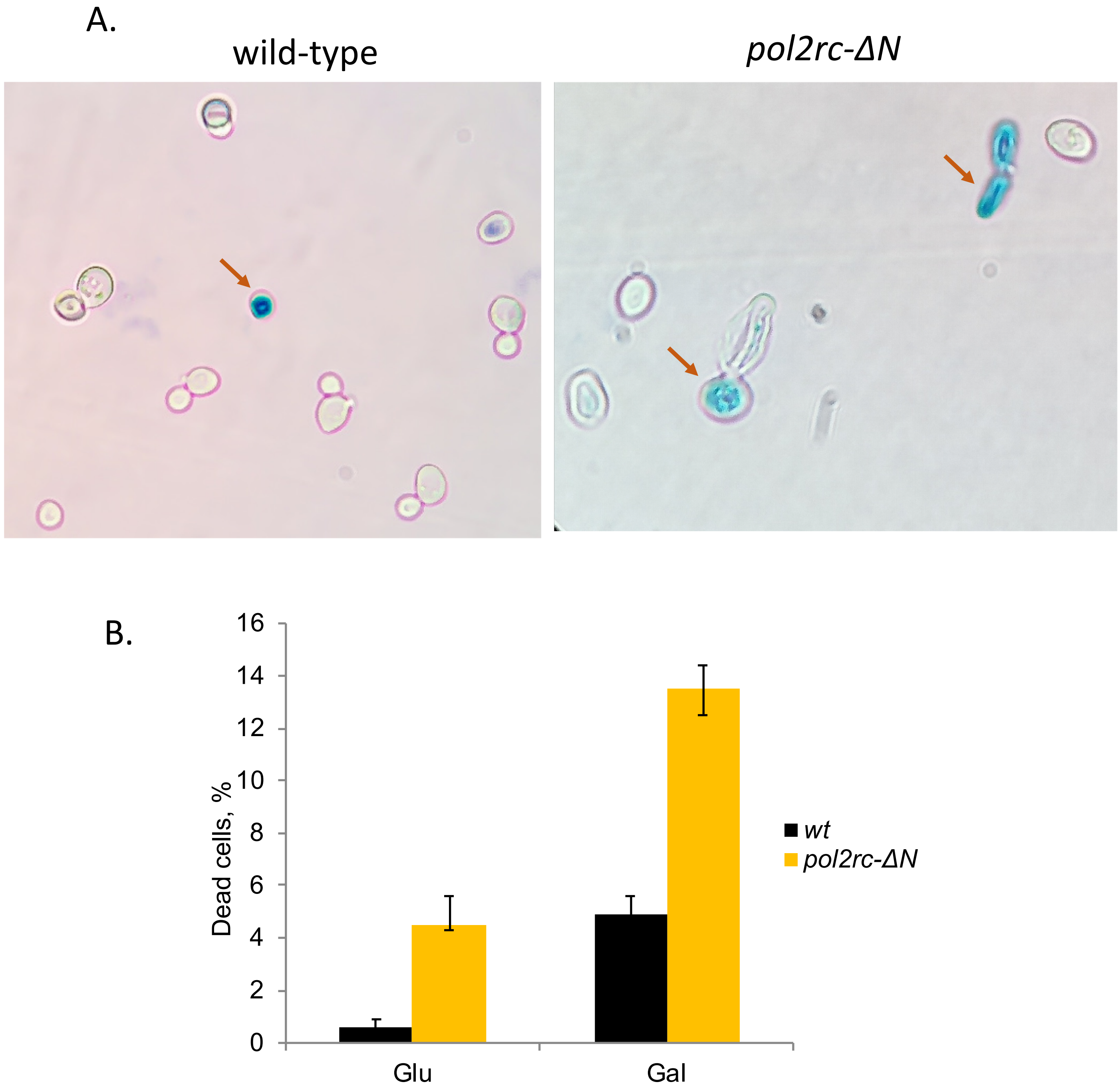


**Supplemental Figure 6. A higher proportion of methylene blue-stained cells in cultures of *pol2rc-ΔN* strains in comparison to wild-type strains.**

A. Example of the appearance of cells stained with methylene blue under microscope, marked by brown arrows.

B. The proportion of the dead cells in wild-type (black) and *pol2rc-ΔN* cells (yellow)

Glu – YPD*, Gal – YPRG*. For each variant, 5000 cells were evaluated. Error bars represent 95% confidence limits.
