## Supplemental results, figures and Tables for "Compensation for the absence of the catalytically active half of DNA polymerase ε in yeast by positively selected mutations in *CDC28* gene": Suppl Fig 7.pdf

*POL2*

*pol2rc-ΔN*

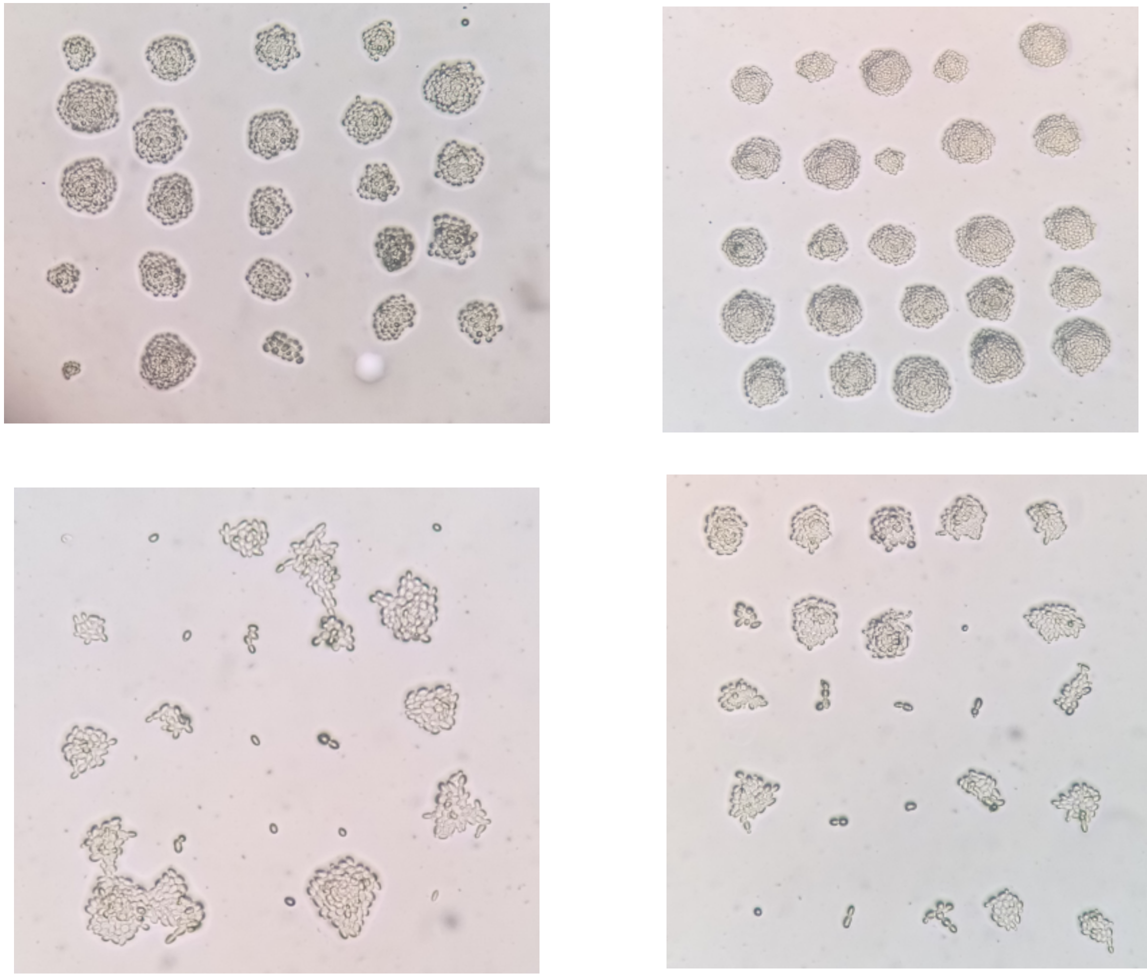

**Supplemental Figure 7. Many cells with *pol2rc-ΔN* have difficulty to start dividing*.***

Photographs of miicrocolonies grown on YPRGAU plates in 24 hours at room temperature (20 $^{o}$C) from single bud-less cells put by micromanipulator in a grid.
