## Supplemental results, figures and Tables for "Compensation for the absence of the catalytically active half of DNA polymerase ε in yeast by positively selected mutations in *CDC28* gene": Suppl Fig 8.pdf

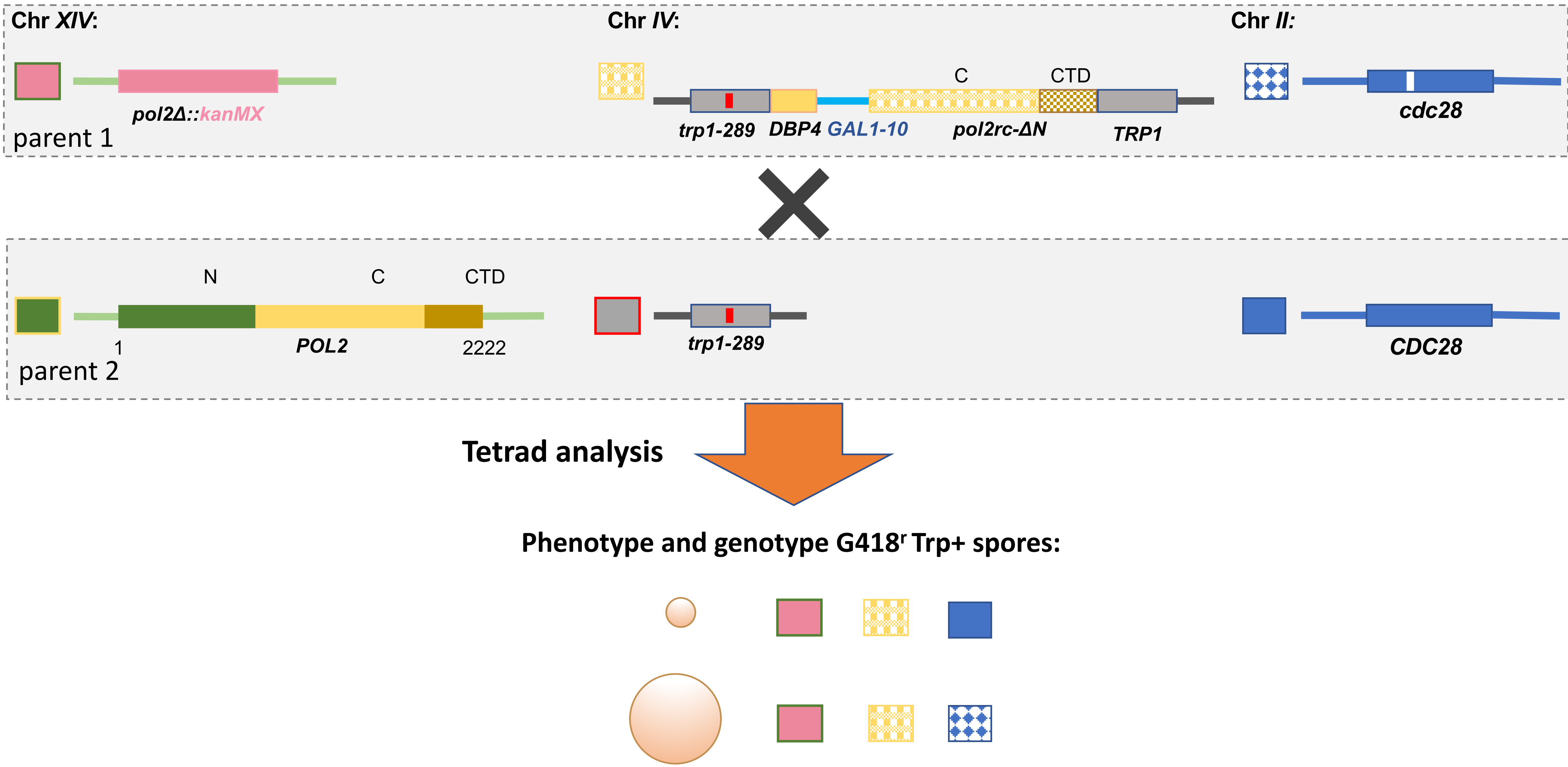


**Supplemental Figure 8. Schematic of analysis to find if single *cdc28* mutations can confer a growth advantage to *pol2rc-ΔN* strains.**
Segregants carrying different *cdc28* mutations were crossed to the wild-type strain, and the *CDC28* gene was sequenced in *POL2 Δ pol2rc-ΔN* isolates. In addition to symbols used in Fig. 1, here we used blue for wild-type *CDC28* and patterned blue for *cdc28* mutations.
