## Supplemental results, figures and Tables for "Compensation for the absence of the catalytically active half of DNA polymerase ε in yeast by positively selected mutations in *CDC28* gene": Suppl Fig 9.pdf

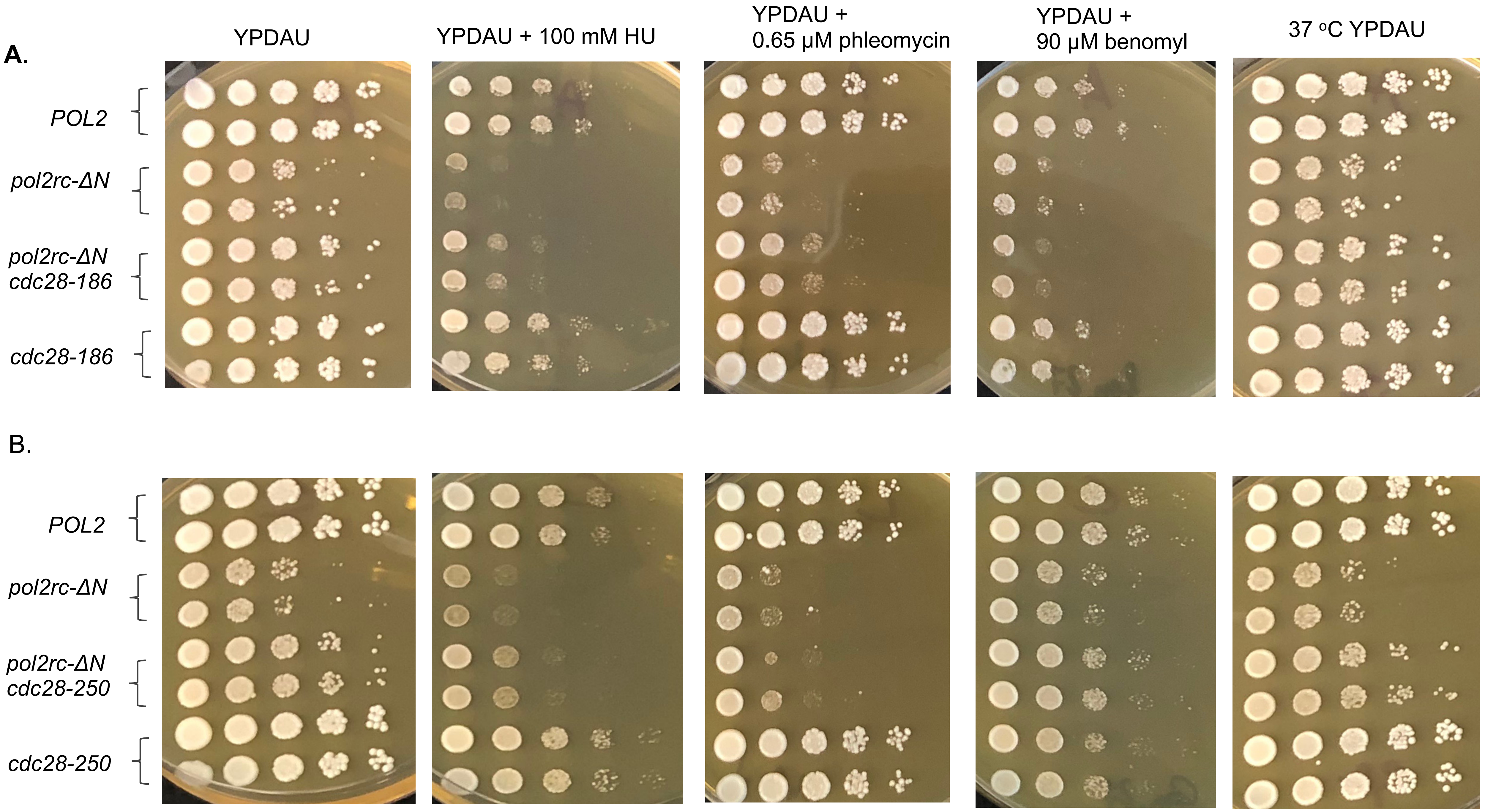

A.
YPDAU
YPDAU + 100 mM HU
YPDAU + 0.65 µM phleomycin
YPDAU + 90 µM benomyl
37 °C YPDAU
POL2
pol2rc-ΔN
pol2rc-ΔN cdc28-186
cdc28-186
B.
POL2
pol2rc-ΔN
pol2rc-ΔN cdc28-250
cdc28-250

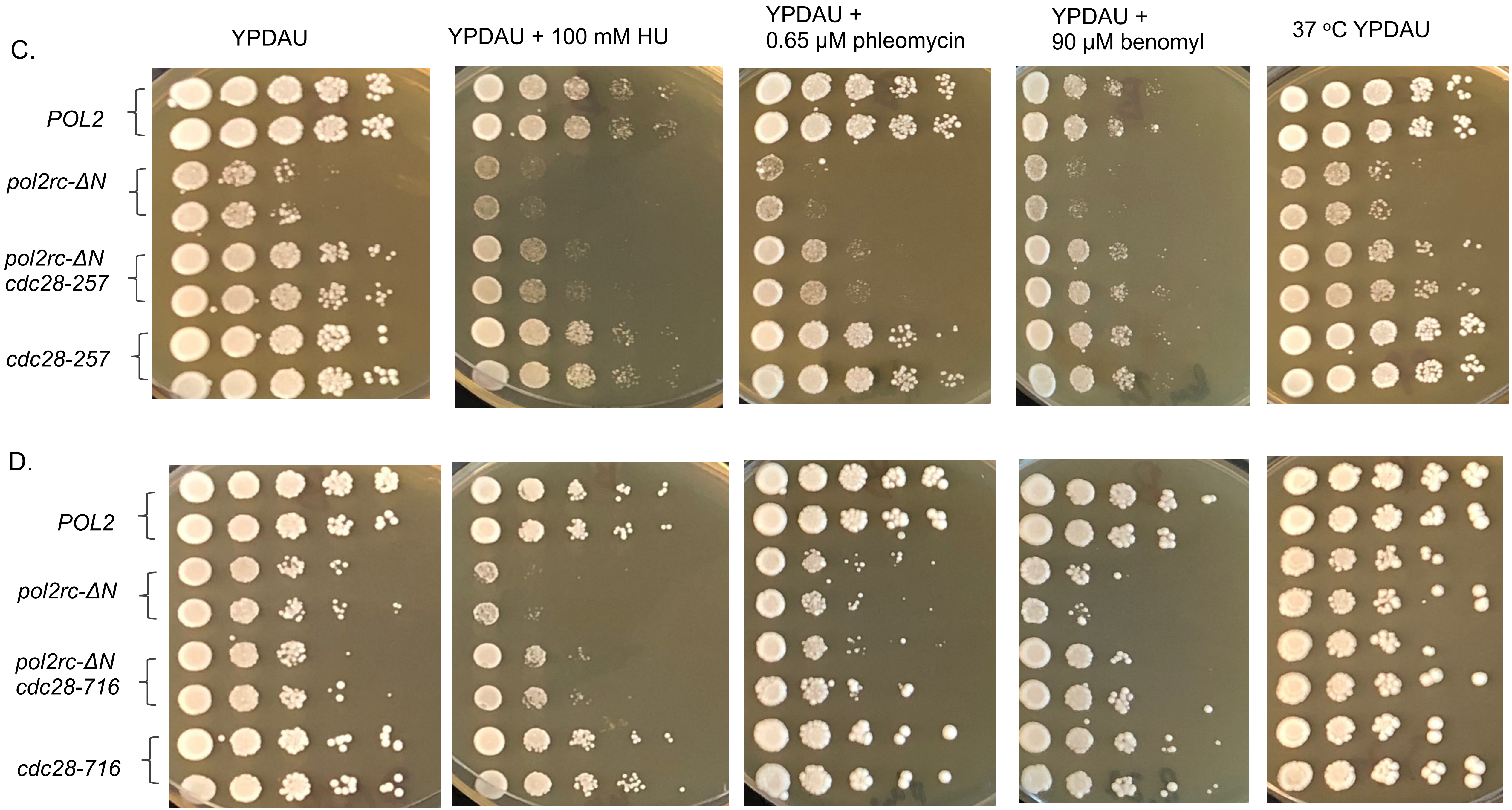


**Supplemental Figure 9. Rescue of growth defects and sensitivity to drugs of *pol2rc-ΔN* by *cdc28* mutations.**
