## Supplemental results, figures and Tables for "Compensation for the absence of the catalytically active half of DNA polymerase ε in yeast by positively selected mutations in *CDC28* gene": Suppl Fig 10.pdf

```
Saccharomyces cerevisiae          --------------msgelanYkrlEKvGEGTYGVVYKAlDlRpGqGqrVVALKKIRLES
Homo sapiens                      ------------------MEDYtKIEKIGEGTYGVVYKgRhKtT---GQVVAmKKIRLES
Caenorhabditis elegans            mdpiregevahegdsVytlnDftKlEKIGEGTYGVVYKgKnRRT---namVAmKKIRLES
Schizosaccharomyces octosporus    ------------------MdnYvKVEKIGEGTYGVVYKAKhKss---GrIVAmKKIRLEd
Candida pseudohaemulonis          ---------------MvelsDfqrqEKIGEGTYGVVYKAlDtRh--nnrVVALKKIRLES
Spizellomyces punctatus           ------------------MdkYdKIEKvGEGTYGVVYKARDRhs---GEIVALKKIRLEt
Apiotrichum porosum               ----------------mSlqnYtKlEKvGEGTYGVVYKAvDlQs---GnfVALKKIRLEa
Cryptococcus amylolentus          ----------------mSldnYhKlEKvGEGTYGVVYKAKDinT---GsIVALKKIRLEa
Kwoniella pini                    ----------------mSldnYtKlEKvGEGTYGVVYKARDlsT---GnfVALKKIRLEa
Malassezia pachydermatis          ------------------MEnYqKIEKvGEGTYGVVYKARDltpGanGrLVALKKIRLEa
Pseudomicrostroma glucosiphilum   ------------------MEnYqKlEKvGEGTYGVVYKARDltaegGGrIVALKKIRLEa
Nematostella vectensis            ------------------MdDfsKIEKIGEGTYGVVYKAKnlKT---GgfaALKKIRLEv
Trichoplax adhaerens              ------------------MdkYlKIEKIGEGTYGVVYKgKnRnT---qQLVALKKIRLEn
Hemicentrotus pulcherrimus        ------------------MEDftKIEKlGEGTYGVVYKgRhKRT---GkIVALKKIRLES
Halocynthia roretzi               ------------------MEDYiKIEKIGEGTYGVVYKgRnKKT---nQyVALKKIRLES
Oryzias luzonensis                ------------------MEDYvKIEKIGEGTYGVVYKgRhKsT---GQVVAmKKIRLES
Rana dybowskii                    ------------------MdeYaKIEKIGEGTYGVVYKgvhKaT---GQIVAmKKIRLEn
Gallus gallus                     ------------------MEDYtKIEKIGEGTYGVVYKgRhKtT---GQVVAmKKIRLES
Drosophila melanogaster           ------------------MEDfeKIEKIGEGTYGVVYKgRnRlT---GQIVAmKKIRLES
Ixodes scapularis                 ------------------MEDYtKVEKIGEGTYGVVYKgKnKKs---GQIVAmKKIRLES

                                                 *L62                    L84* *L86
Saccharomyces cerevisiae          EDEGVPSTAIREISLLKEL-Kdd----NIVrLyDIvHsdaHKLYLVFEFLDlDLKrYMeg
Homo sapiens                      EeEGVPSTAIREISLLKEL-RHP----NIVsLqDvLmQdS-rLYLiFEFLsMDLKKYlDS
Caenorhabditis elegans            EDEGVPSTAvREISLLKEL-qHP----NvVgLeavimQEn-rLfLiFEFLsfDLKrYMDq
Schizosaccharomyces octosporus    EsEGVPSTAIREISLLKEv-ndesnrsNcVrLlDILHaES-KLYLVFEFLDMDLKKYMDr
Candida pseudohaemulonis          EDEGVPSTAIREISLLKEm-Rdd----NIVrLyDIiHsdSHKLYLVFEFLDlDLKKYMel
Spizellomyces punctatus           EDEGVPSTAIREISLLKEL-KHP----NIVrLlDIvHnda-KLYLiFEFLDlDLKKYMDT
Apiotrichum porosum               EDEGVPSTsIREISLLKELSKdd----NIVkLlDIvHQda-KLYLVFEFLDlDLKrYMDS
Cryptococcus amylolentus          EDEGVPSTsIREISLLKELSKdd----NIVkLlDIvHsEa-KLYLVFEFLDMDLKKYMDS
Kwoniella pini                    EDEGVPSTsIREISLLKELSKdd----NIVkLlDIvHsda-KLYLVmEFLDMDLKKYMDN
Malassezia pachydermatis          EDEGVPSTAIREISLLKEL-Rde----NIVrLyeIiHQES-rLYLVFEFLDlDLKKYMDN
Pseudomicrostroma glucosiphilum   EDEGVPSTAIREISLLKEL-Knd----NIVrLyDIiHQES-KLYLVFEFLDlDLKKYMDN
Nematostella vectensis            EDEGiPSTAvREISLLKELrhHP----NvVeLqhILHQEp-KLYLVFEyLtcDLKKhlDT
Trichoplax adhaerens              EeEGiPSTAIREvSLLKEL-KHP----NIVdLievLYEES-KLYLVFEFLDMDLKrYlDT
Hemicentrotus pulcherrimus        EeEGVPSTAIREISLLKEL-yHP----NIVlLeDvLmEpn-rLYLVFEyLtMDLKKYMeS
Halocynthia roretzi               EeEGVPSTAIREISiLKEL-qHP----NIVsLlDvLlQES-KLYLVFEFLqMDLKKYMDS
Oryzias luzonensis                EeEGVPSTAvREvSLLqEL-KHP----NvVrLlDvLmQES-rLYLiFEFLsMDLKKYlDS
Rana dybowskii                    EeEGVPSTAIREISLLKEL-qHP----NIVcLlDvLmQdS-rLYLiFEFLsMDLKKYlDS
Gallus gallus                     EeEGVPSTAIREISLLKEL-hHP----NIVcLqDvLmQda-rLYLiFEFLsMDLKKYlDT
Drosophila melanogaster           dDEGVPSTAIREISLLKEL-KHe----NIVcLeDvLmEEn-riYLiFEFLsMDLKKYMDS
Ixodes scapularis                 EDdGVPSTAIREItLLKEL-nHr----NIVrLqDvimQEn-KvYLVFEFLsMDLKKhlDT

Saccharomyces cerevisiae          IPk-d-QpLgadIVKkfMmQlcKGIayCHShRiLHRDLKPQNLLInKdGnlKLgDFGLAR
Homo sapiens                      IPp-G-QymDssLVKSYLYQILQGIvFCHSRRVLHRDLKPQNLLIDdKGtIKLADFGLAR
Caenorhabditis elegans            Lgk-d-EyLpletlKSYtfQILQamcFCHqRRViHRDLKPQNLLvDnnGaIKLADFGLAR
Schizosaccharomyces octosporus    IsetGAsaLDPrLVqkfaYQlvnGvnFCHSRRiiHRDLKPQNLLIDKEGnlKLADFGLAR
Candida pseudohaemulonis          IPq-G-agLEPsMVKrfMhQlIKGIkhCHShRVLHRDLKPQNLLIDKdGnlKLADFGLAR
Spizellomyces punctatus           qsn----gLsapLiKSYMYQlIKGIhyCHchRiLHRDLKPQNLLIDqQGmlKLADFGLAR
Apiotrichum porosum               LPd-k-EaLpPnLVKkftYQlIKGlfyCHvhRVLHRDLKPQNLLIDKEGnlKLADFGLAR
Cryptococcus amylolentus          Lge-k-dgLgPdMVKkfsYQlIKGIyFCHghRiLHRDLKPQNLLInKgGdlKiADFGLAR
Kwoniella pini                    Igd-k-dgLgPnMVKkftYQlvKGlyyCHAhRiLHRDLKPQNLLInKEGnlKiADFGLAR
Malassezia pachydermatis          Vad-kpEgLgPeIVmkftYQlvRGIyFCHAhRiLHRDLKPQNLLIDKEGnlKLADFGLAR
Pseudomicrostroma glucosiphilum   Vag-tAdgLgPeIVKkftfQlLRGIyyCHAhRiLHRDLKPQNLLIDKtGnlKLADFGLAR
Nematostella vectensis            trg----mLDktLVKSYLYQItnaIyFCHARRiLHRDLKPQNLLIDsKGlIKLADFGLgR
Trichoplax adhaerens              LPk-G-ktiDamLmKSYLYQILlGvvyCHShRVLHRDLKPQNLLInsKGcIKLADFGLgR
Hemicentrotus pulcherrimus        L-k-G-kqmDPaLVKSYLhQmvdGIlFCHSRRiLHRDLKPQNLLIDnnGtIKLADFGLAR
Halocynthia roretzi               IPa-G-kymDkeLVKSYtYQILQGItFCHSRRVLHRDLKPQNLLIDKnGiIKLADFGLAR
Oryzias luzonensis                IPs-G-QymDPmLVKSYLYQILEGIyFCHrRRVLHRDLKPQNLLIDnKGvIKLADFGLAR
Rana dybowskii                    IPs-G-QyLEamLVKSYLYQILQGIiFCHARRVLHRDLKPQNLLIDsKGvIKLADFGLAR
Gallus gallus                     IPs-G-QyLDrsrVKSYLYQILQGIvFCHSRRVLHRDLKPQNLLIDdKGvIKLADFGLAR
Drosophila melanogaster           LPv-d-khmEseLVrSYLYQItsaIlFCHrRRVLHRDLKPQNLLIDKsGlIKvADFGLgR
Ixodes scapularis                 LPk-n-QsmDtktVKSYLkQILEGIlFCHrRRVLHRDLKPQNLLIDqKGnIKLADFGLAR
```

```
Saccharomyces cerevisiae             AFGvPLRaYTHEiVTLWYRAPEVLLGgkqYSTgVDtWSIGCIFAEMcnrKPiFsGDSEID
Homo sapiens                         AFGIPiRVYTHEVVTLWYRsPEVLLGSaRYSTPVDiWSIGtIFAElATKKPLFHGDSEID
Caenorhabditis elegans               AiGIPiRVYTHEVVTLWYRAPEiLmGaqRYSmgVDMWSIGCIFAEMATKKPLFqGDSEID
Schizosaccharomyces octosporus       sFGvPLRnYTHEiVTLWYRAPEVLLGSrhYSTgVDiWSVGCIFAEMirrtPLFpGDSEID
Candida pseudohaemulonis             AFGvPLRaYTHEVVTLWYRAPEiLLGgkqYSTgVDMWSVGCIFAEMcnrKPLFpGDSEID
Spizellomyces punctatus              AFGvPLRtYTHEVVTLWYRAPEiLLGSkhYSTaVDMWSVGCIFAEMclrhPLFpGDSEID
Apiotrichum porosum                  AFGIPLRtYTHEVVTLWYRAPEVLLGSrhYSTaiDMWSVGCIFAEMAmrsPLFpGDSEID
Cryptococcus amylolentus             AFGIPLRtYTHEVVTLWYRAPEVLLGSrhYSTaiDMWSVGCIvAEMATrqPLFpGDSEID
Kwoniella pini                       AFGIPLRtYTHEVVTLWYRAPEVLLGSrhYSTaiDMWSVGCIyAEMAmrqPLFpGDSEID
Malassezia pachydermatis             AFGIPLRtYTHEVVTLWYRAPEVLLGSrhYnTaiDMWSVGCIFAEMAmrtPLFpGDSEID
Pseudomicrostroma glucosiphilum      AFGIPLRtYTHEVVTLWYRAPEVLLGSrhYSTaiDMWSVGCIFAEMAmKsPLFpGDSEID
Nematostella vectensis               AFGIPvRaYTHEVVTLWYRAPEVLLGgqRYScPiDvWSIGtIFAEMvTKRPLFHGDSEID
Trichoplax adhaerens                 AFGvPvRVYTHEVVTLWYRAPEVLLGStRYScPlDiWStGtIFAEMwlrRPLFqGDSEID
Hemicentrotus pulcherrimus           AFGIPvRVYTHEVVTLWYRAPEVLLGStRYacPiDMWSlGCIFAEMvTKRPLFHGDSEID
Halocynthia roretzi                  AFGIPvRVYTHEVVTLWYRAPEVLLGasRYSTPVDiWSIGtIFAEMATKKPLFHGDSEID
Oryzias luzonensis                   AFGvPvRVYTHEVVTLWYRAPEVLLGSpRYSTPVDvWStGtIFAElATKKPLFHGDSEID
Rana dybowskii                       AFGIPvRVYTHEVVTLWYRAPEVLLGSvRYSTPVDvWSIGtIFAEiASKKPLFHGDSEID
Gallus gallus                        AFGIPvRVYTHEVVTLWYRsPEVLLGSalYSTPVDiWSIGtIFAElATKKPLFHGDSEID
Drosophila melanogaster              sFGIPvRIYTHEiVTLWYRAPEVLLGSpRYScPVDiWSIGCIFAEMATrKPLFqGDSEID
Ixodes scapularis                    AFGIPiRVYTHEiVTLWYRAPEVLLGSpRYSTPiDiWSIaCIFvEMinKRPLFHGDSEID

                                                        *I239
Saccharomyces cerevisiae             qiFkIFRvLGTPNEaIWPdivyLPDfKpsFPqWRrkdLsqvVps-------LDprGIDLL
Homo sapiens                         qLFRIFRaLGTPNnEVWPeVeSLqDYKNTFPKWKpgsLashVKN-------LDENGLDLL
Caenorhabditis elegans               ELFRIFRvLGTPtEleWnGVeSLPDYKaTFPKWRenfLrdkfydkktgkhlLDDtafsLL
Schizosaccharomyces octosporus       EiFkIFqvLGTPNEEVWPGVTlLqDYKSTFPRWKrvdLhrvIpN-------geEaaIeLL
Candida pseudohaemulonis             EiFRIFRiLGTPNEEVWPdVTyLPDfKtTwPKWErsdLgphIps-------LDrdGvDLL
Spizellomyces punctatus              EiFRIFRTLGTPNEEIWPnVTtLPDYKenFPiWtaqnLakvlpN-------LeseGvDLL
Apiotrichum porosum                  EiFRIFRlLGTPdEEsWPGirSLPDYKSsFPqWhqaeLggaVsg-------LDaNGLDLL
Cryptococcus amylolentus             EiFRIFRvLGTPdEdVWPGVrSLPDYKpTFPqWspinLgdvVKg-------LDadGMDLv
Kwoniella pini                       EiFRIFRvLGTPdEdVWPGVraLPDYKpTFPqWnaveLksaVKg-------LDENGsDLL
Malassezia pachydermatis             EiFRIFRTLGTPNdEVWPGVqSLPDYKtTFPKWngvpLksaVsg-------LDDtGLDLL
Pseudomicrostroma glucosiphilum      EiFRIFRTLGTPtdaVWPGVktLPDYKasFPqWsgmpLakaVpN-------LDaNGIDLL
Nematostella vectensis               qLFRIFRiLGTPtEEtWkGVTSLPDYKpTFPKWagdgLkkaVpq-------LDsdGLDLL
Trichoplax adhaerens                 ELFRIFRiLGTPdddIWPGVSSLPefKSsFPKWskqsydtfVpN-------msEsGIDLL
Hemicentrotus pulcherrimus           qLFRIFRTLGTPtdEIWPGVTqLqDYKSTFPmWtkpnikgaVKg-------mDEeGLDLL
Halocynthia roretzi                  qLFRIFRvLGTatEddWPGVTSLkDYKrTFPKWKkgmvvesVKN-------LnEeGIDLL
Oryzias luzonensis                   qLFRIFRTLGTPNndVWPdVeSLPDYKNTFPKWKggsLssmVKN-------LDKNGLDLL
Rana dybowskii                       qLFRIselwGTPNnEVWPeVeSLqDYKNTFPKWKggsLaanVKN-------iDKeGLDLL
Gallus gallus                        qLFRIFRaLGTPNndVWPdVeSLqDYKNTFPKWKpgsLgthVqN-------LDEdGLDLL
Drosophila melanogaster              qLFRmFRiLkTPtEdIWPGVTSLPDYKNTFPcWstnqLtnqlKN-------LDaNGIDLi
Ixodes scapularis                    qLFRIFRTLGTPtEdtWPGVTkLPDYKSsFPnWseniLrsllKN-------mDDdGIDLL

Saccharomyces cerevisiae             dKlLaYDPinRISArRAaiHPYFqEs--------------------
Homo sapiens                         SKMLIYDPAkRISgKmALnHPYFNDlDnqikkm-------------
Caenorhabditis elegans               eglLIYDPslRlnAKKALvHPYFDnmDtS---kLPagnyrGeLelf
Schizosaccharomyces octosporus       SaMLVYDPAhRISAKRALqqPYlrEygeTfv---------------
Candida pseudohaemulonis             eqMLnYDPsnRISAKRALvHPYFqEdNdeaydsyPrsvnmG-----
Spizellomyces punctatus              SRlLVYDPAqRlSAKRALsHPYFaDvDtSksl--------------
Apiotrichum porosum                  AqtLIYDPAqRISAKRALqHPYFatsvaAaangta-----------
Cryptococcus amylolentus             AqtLVfDPAhRISAKRALrHPYFDtvNlAaa---------------
Kwoniella pini                       AqtLIfDPAhRISAKRALqHPYFtstypA-----------------
Malassezia pachydermatis             hhMLIYDPAiRISAKRALqHPYFasvsaA-----------------
Pseudomicrostroma glucosiphilum      kgMLVYDPAgRvSAKRgLnHaYFqslpgSamas-------------
Nematostella vectensis               kKMLIYDPAlRISAKtsLkHPYFlndpkfdinsLPktpevdsVm--
Trichoplax adhaerens                 SKMLIYDPAnRISgKRALsHPYFDDlDkS---tLPtdrws------
Hemicentrotus pulcherrimus           eKMLIYDPAkRItAKasmrHPYFDnipdlsdrlqPirs--------
Halocynthia roretzi                  qKcLVYDPAkRISAKaALmHPYFNnlDkklclvLtsalfhe-----
Oryzias luzonensis                   AKMLIYnPpkRISAreAmtHPYFDDlDkS---tLPaacinGv----
Rana dybowskii                       AKMLVYDPAkRISArKALlHPYFDDlDkS---sLPanqirn-----
Gallus gallus                        SKMLIYDPAkRISgKmALnHPYFDDlDkS---tLPanlikkf----
Drosophila melanogaster              qKMLIYDPvhRISAKdiLeHPYFNgfqsglvrn-------------
Ixodes scapularis                    eKMLVYDPvrRISAKdcLdHPYlND---------------------
```

**Supplementary Figure 10. Multiple alignment of CMGC/CDK/CDC2 protein kinase orthologs.** Blue capital letters stand for highly conserved positions, as suggested by the EBI web server (https://www.ebi.ac.uk/Tools/msa/muscle/). Asterisks show positions of amino acid changed in *cdc28* mutants found in our work (labeled either on the right of left to the asterisk). Protein identifiers: NP_001307847.1 (*Homo sapiens*), NP_009718.3 (*Saccharomyces cerevisiae*),

XP_016605543.1 (*Spizellomyces punctatus*), XP_017992564.1 (*Malassezia pachydermatis*), XP_028477809.1 (*Apiotrichum porosum*), XP_025347166.1 (*Pseudomicrostroma glucosiphilum*), XP_019012246.1 (*Kwoniella pini*), XP_018995495.1 (*Cryptococcus amylolentus*), XP_024712237.1 (*Candida pseudohaemulonis*), XP_013017915.1 (*Schizosaccharomyces octosporus*), P13863 (*Gallus gallus)*, P34556 (*Caenorhabditis elegans*), P23572 (*Drosophila melanogaster*), BAC98412.1 (*Halocynthia roretzi*), Q9DG98 (*Oryzias luzonensis*), Q9W739 (*Rana dybowskii*), BAA23218.1 (*Hemicentrotus pulcherrimus*), XP_001635734.1 (*Nematostella vectensis*), XP_002111844.1 (*Trichoplax adhaerens*), XP_002415172.1 (*Ixodes scapularis*).
