## Supplemental results, figures and Tables for "Compensation for the absence of the catalytically active half of DNA polymerase ε in yeast by positively selected mutations in *CDC28* gene": Suppl Table 1.docx

**Supplemental Table 1**. **Mutator effect of *pol2rc-ΔN* is reduced 60% in strain without the catalytic subunit of pol ζ, Rev3.**

| Parent diploid and type of segregants | Rate of Can^r^ x 10^-7^  (95% confidence limits) |
| --- | --- |
| YEE303r3hΒ/+:  *POL2rc* *rev3*::*hph* segregants (n=2)* | 1.5  (0.9-3.3) |
| YEE304r3hΒ/+:  *pol2rc-ΔN rev3::hph* segregants (n=9) | 32  (15-50) |
| YEE304r3hΒ/+:  *pol2rc-ΔN REV3* segregants (n=9) | 83  (61-148) |

*- for each segregant, 9-18 independent cultures were used to estimate mutation rates
