## Supplemental results, figures and Tables for "Compensation for the absence of the catalytically active half of DNA polymerase ε in yeast by positively selected mutations in *CDC28* gene": Suppl Table 2.docx

**Supplemental Table 2.** **Differences in mutant frequencies with or without UV-light in *pol2rc-ΔN* *vs.* wild-type strains***

| UV dose | Mutant frequency × 10^-7^, mean ± SEM | | | | Survival %,  mean ± SEM | |
| --- | --- | --- | --- | --- | --- | --- |
|  | Can^r^ | | His^+^ | |  |  |
|  | wt | *pol2-ΔN* | wt | *pol2-ΔN* | wt | *pol2-ΔN* |
| No UV | **32.7** ± 10.8 | **236.6** ± 33.7  p<0.00001 | **0.3** ± 0.08 | **2.3** ± 0.4  p<0.00001 | **100** | **100** |
| UV, 10 J/m^2^ | **283.8** ± 54.7 | **453.7** ± 111.8  p=0.09029 | **6.0** ± 0.9 | **10.7** ± 1.1  p= 0.001684 | **93.7** ± 4.6 | **94.8** ± 5.9  p= 0.459157 |
| UV, 20 J/m^2^ | **860.9** ± 203.2 | **697.7** ± 150.4  p= 0.261212 | **16.5** ± 1.4 | **25.2** ± 1.9  p=0.000298 | **77.3** ± 2.6 | **73.9** ± 4.9  p=0.269633 |
| UV, 40 J/m^2^ | **2602.3** ± 684.5 | **1393.5** ± 285.3  p= 0.05576 | **59.2** ± 2.9 | **72.7** ± 3.9  p=0.004302 | **43.5** ± 2.0 | **35.3** ± 2.0  p= 0.002848 |

* The P values for statistically significant differences are in red.
