## Supplemental results, figures and Tables for "Compensation for the absence of the catalytically active half of DNA polymerase ε in yeast by positively selected mutations in *CDC28* gene": Suppl Table 3.docx

**Supplemental Table 3. The *pol2rc-ΔN* elevated chromosome instability measured by the α-test (illegitimate mating).**

| Strain | Frequency of illegitimate mating x10^-7^ (95% confidence limits) | | | | | |
| --- | --- | --- | --- | --- | --- | --- |
|  | Illegitimate hybridization | Point mutations and primary lesions in *MATα* | Chromosome *III* loss | Loss of Chr. *III* right arm | Conversion *HMRa*  ->*MATα* | Recombination between *HMRa*  and *MATα* |
| wild-type | **35.6**  (20.3-42.8) | **29.6**  (16.8-35.5) | **2.2**  (1.3-2.7) | **2.5**  (1.4-3.1) | **0.2**  (0.1 - 0.22) | **0.15**  (0.09 - 0.19) |
| *pol2rc-ΔN* | **256.6**  (132.7 - 350.5) | **126.3**  (65.3 - 172.5) | **47.6**  (24.6-65.0) | **73.7**  (38.1 - 100.6) | **0.7**  (0.4 - 0.9) | **0.3**  (0.1 - 0.4) |
