## Supplemental results, figures and Tables for "Compensation for the absence of the catalytically active half of DNA polymerase ε in yeast by positively selected mutations in *CDC28* gene": Suppl Table 6.docx

**Supplemental Table 6. Types of mutations found in genomes of *pol2rc-ΔΝ* strains.**

| Type of substitution | Amount of mutations found in all genomes | | Amount of independent mutations (taking into account common origin of sequenced genomes) | |
| --- | --- | --- | --- | --- |
|  | Number of mutations | % | Number of independent mutations | % |
| GC->CG | 116 | 39.7 | 89 | 45.2 |
| AT->TA | 50 | 17.1 | 39 | 19.8 |
| AT->CG | 12 | 4.1 | 12 | 6.1 |
| AT->GC | 35 | 12.0 | 25 | 12.7 |
| GC->AT | 28 | 9.6 | 15 | 7.6 |
| GC->TA | 49 | 16.8 | 16 | 8.1 |
| Complex (AAA->TAT) | 2 | 0.7 | 1 | 0.5 |
| Total | 292 | 100 | 197 | 100 |
