## Supplemental results, figures and Tables for "Compensation for the absence of the catalytically active half of DNA polymerase ε in yeast by positively selected mutations in *CDC28* gene": Supplemental Results 1.docx

***Estimation of the probabilities of recurrent mutations in the same gene.***

Let us consider a random 28 × 6075 matrix (the rows of the matrix represent independent sequenced genomes, the columns represent number of functional yeast genes used in this study). If there is a mutation in the gene *j* from the genome *i*, *aij* = 1, otherwise *aij* = 0. Thus, the entries *aij* are independent variables taking the value 1 with the probability *p* and the value 0 with the probability 1-*p*. Note that the probability is the same for all *i* and *j*. Our goal is to estimate the probability P_≥5_ that there is a gene with at least 5 mutations (the total number of mutations found in 28 independent genomes is 197, **Table 1**). Let us first estimate *p*. Since the variables are independent, the expected value of the total number of mutations is

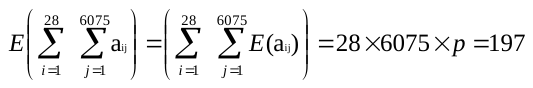

whence

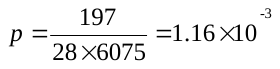

Now let us calculate the probability of at least 5 mutations in a fixed gene *j*. The probability of exactly *k* mutations in this gene is

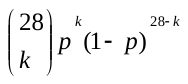
.

Thus, the probability of at most 4 mutations is

*p*_≤4_ =
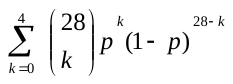

and the probability of at least 5 mutations is

*p*_≥5_ = 1 – *p*_≤4_ ≈ 2 × 10^-10^.

Finally, since the events “at least five mutations in the *j*-th gene" are independent for different *j*, the probability P_≥5_ is

P_≥5_ = 1 – (1 – *p*_≥5_)^6075^ = 1.2 × 10^-6^.

The same logic was applied to 3 and 4 multiple independent mutations.

3 mutations, P = 0.0302

4 mutations, P = 0.0002

5 mutations, P = 1.2 × 10^-6^

Thus, the probability to observe 3 independent mutations in the same gene is marginally significant whereas 4 mutations are highly significant.

We also performed a simulation experiment as a control, we randomly populated the 28 × 6075 matrix with 197 mutations and estimated a weight

W_random_ =
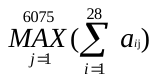

We repeated this procedure 1000 times and did not find any simulated matrix with the weight W_random_ ≥ 5. This suggested that the probability to observe 5 or more mutations in a gene is less than 0.001. This is consistent with our analytical estimates (P_≥5_  ≈ 1.2 × 10^-6^).

Thus, the probability to find 5 or more mutations in a gene is extremely small suggesting that this event is likely to be biologically important.
